# Dapagliflozin mitigates hypoxia-induced metabolic stress, kidney tubular cell death and fibrosis

**DOI:** 10.64898/2025.12.13.693777

**Authors:** Akihiro Minakawa, Jamal El Saghir, Céline C. Berthier, Moritz Lassé, Soumya Maity, Judy Baek, Damian Fermin, Anja Billing, Chenchen He, Felix Eichinger, John Hartman, Viji Nair, Matthew Fischer, Virginia Vega-Warner, Fadhl Alakwaa, Bradley Godfrey, Phillip J. McCown, Rajasree Menon, Fatih Demir, Subramaniam Pennathur, Kumar Sharma, Jennifer A. Schaub, Markus M. Rinschen, Matthias Kretzler, Jennifer L. Harder

**Author notes:** Authors contributed equally (Akihiro Minakawa, Jamal El Saghir, Céline C. Berthier).

## Abstract

Sodium-glucose co-transporter 2 inhibitors (SGLT2-i) slow progression of kidney disease but therapeutic mechanisms remain elusive. Here we report the beneficial effect of dapagliflozin on hypoxia-mediated kidney tubular epithelial cell injury, a contributing factor to kidney disease progression, using a human pluripotent stem cell (hPSC)-derived hypoxic kidney organoid model. Hypoxic organoids showed increased expression of Hypoxia Inducible Factor (HIF)-associated transcriptional targets, decreased tricarboxylic acid (TCA) cycle metabolites and mitochondrial β-oxidation protein expression, and activated unfolded protein response. A transcriptional signature derived from hypoxic organoids 1) identified a subgroup of individuals whose kidney disease subsequently progressed, and 2) correlated with worse tubular injury. Dapagliflozin enhanced mitochondrial stress response resulting in reversed hypoxia-induced tubular epithelial cell apoptosis, reactive oxygen species (ROS) accumulation, and organoid fibrosis. These results indicate that dapagliflozin may contribute to improved kidney disease outcomes by attenuating hypoxia-induced metabolic stress-mediated tubular epithelial cell injury.

## INTRODUCTION

Kidney disease affects more than 10% of the population and is the 3rd fastest growing cause of death worldwide^1,2^. However, effective treatments to halt or reverse disease course have been slow to develop. This reflects limited *in vitro* modeling success due to myriad factors including complex kidney physiology and architecture with more than 51 main kidney cell types as well as human specificity of kidney gene expression^3–5^. A landmark breakthrough in reduction in the risk for chronic kidney disease (CKD) progression resulted from the introduction of SGLT2-i^6,7^. Beyond increased tubuloglomerular feedback, decreased proximal tubular oxygen usage via decreased energetic demand, and enhanced nutrient deprivation signaling and autophagy, SGLT2-i are proposed to promote oxygen deprivation signaling and contribute to tubular repair^8–10^.

Intriguingly, the risk of acute kidney injury (AKI) was also noted to be lower with SGLT2-i treatment, even with use in lower kidney perfusion states such as heart failure and mild hypovolemia induced by SGLT2-i use as well as contrast-induced arteriolar constriction^11–14^. Acute hypoperfusion injury with resulting tissue oxygen deprivation is a frequent mechanism for acute tubular injury; hypoxia is also proposed to be a significant contributing factor to development and progression of CKD via promotion of inflammation and fibrosis^15–18^. Correlation between degree of hypoxia and tubulointerstitial injury was also observed in study of heavily proteinuric rat models^19^. Indeed, proteinuria is an independent risk factor for AKI^20–22^; a proposed contributing mechanism stems from increased oxygen utilization by proximal tubular cells due to increased energetic demands related to uptake and catabolism of filtered proteins^23^. Multiple studies have demonstrated a protective effect of SGLT-i in proteinuric kidney disease^6,7,24^. However, experimentally exploring SLGT2-i’s impact on hypoxia-induced changes is difficult to discern with *in vivo* models due to complex kidney physiology.

In this study, we investigated whether SGLT2 inhibition protects kidney tubular epithelial cells from hypoxia-mediated injury. Using a hypoxic kidney organoid model generated from human pluripotent stem cells (hPSCs), we demonstrate the relevance of hypoxia-related gene expression to human kidney disease as well as the modulating effects of dapagliflozin on tubular epithelial cell energetics and injury, experimentally supporting observations in humans with AKI.

## RESULTS

### hPSC-kidney organoids exhibit expected transcriptional and metabolic hypoxic response

To model transient ischemia of kidney tissue, we generated kidney organoids from hPSCs and exposed them to hypoxic conditions for 24 hours (Figures 1A, S1A). Nuclear HIF-1α accumulated in hypoxic organoid tubular cells (Figure 1B) and expression of known transcriptional targets of HIF-1 (including *SLC2A1*, *ELGN1*, *VEGFA* and its protein VEGF-A)^23^ significantly increased under hypoxic conditions (Figure 1C-D). These results were confirmed in organoids generated from two additional hPSC lines (Figure S1B-C). Further, GLUT-1 localization to tubular cell membrane increased while glucose level in media was demonstrably decreased under hypoxic conditions consistent with enhanced glucose utilization (Figure 1E). Transcriptional expression of genes involved in glucose metabolism, sirtuin and HIF-1 signaling were increased while genes involved in mitochondrial function were decreased following hypoxia (Figure 1F-G).

**Figure 1.**
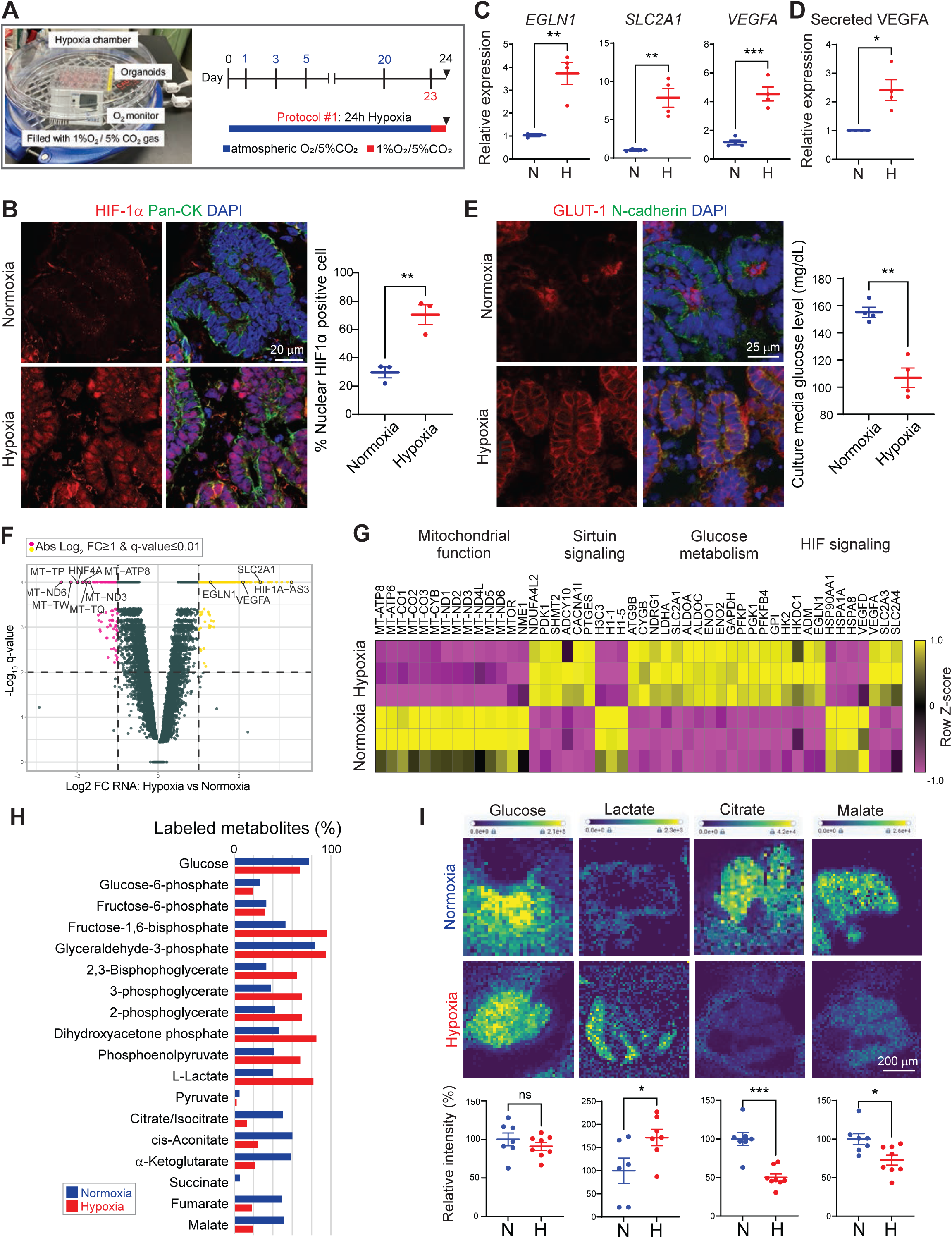
hPSC-kidney organoids exhibit expected transcriptional and metabolic hypoxic response. **(A)** Schematic (left) for hypoxia chamber housing kidney organoids with timeline (right) for 24h Hypoxia Protocol #1. **(B)** Immunofluorescence and quantification (n=3) of nuclear HIF-1α expression in organoids under normoxia (top) or hypoxia (bottom). **(C and D)** Expression of *EGLN1*, *SLC2A1* and *VEGFA* transcripts (C) and secreted VEGF protein levels (D) in hypoxic organoids (n=4). **(E)** Immunofluorescence of GLUT-1 expression in normoxic (top) or hypoxic (bottom) organoids. Scatter plot (right) of glucose concentrations in organoid culture media after 24h hypoxia (n=4). **(F)** Volcano plot of DE genes in hypoxic vs. normoxic organoids. DE transcripts meeting the significance threshold are represented by colored dots, with increased (yellow, upper right quadrant) or decreased (pink, upper left quadrant) relative expression. Selected highly DE transcripts are labeled. Y axis maximum adjusted for visualization purposes. **(G)** Heatmap of selected DE genes (FDR≤0.01 & abs Log_2_ FC≥1) from the top 20 Ingenuity canonical pathways (*p*-value<0.05) in hypoxic vs. normoxic organoids. **(H)** Histogram comparing ^13^C-glucose labeled glycolysis and TCA cycle pathway analytes in hypoxic vs. normoxic organoids after 3h exposure (15 organoids pooled each condition). **(I)** MALDI-MSI images and quantification (6-8 organoids total) of glycolysis and TCA pathway analytes in hypoxic and normoxic organoids evaluated in parallel with (H). Yellow, highest signal intensity. Data displayed as mean +/- SEM; n=experimental replicates; Unpaired t-test (B, C, E and I), Welch’s t-test (D); \**p*<0.05, \*\**p*<0.01, \*\*\**p*<0.001, ns, not significant. IF markers: nuclei (DAPI), tubular epithelial membrane (N-cadherin/pan-cytokeratin (pan-CK)); white scale bars, as labeled. **Related data** Figure S1.

Following hypoxia, tubular architecture remained intact with functional mitochondrial as assessed by pan-cytokeratin and ATP5A (Figure S1D) and oxygen consumption normalized post-reoxygenation (Figure S1E). After 2 days of reoxygenation (Figure S1F), hypoxic organoids were indistinguishable from control organoids by culture media glucose levels, transcript levels of HIF-1 target genes as well as fibrosis-related genes *TGFB1* and *COL1A1*, architecture of tubules and podocytes (N-cadherin, pan-cytokeratin and podocalyxin), and by return of HIF-1 and GLUT-1 expression to normoxic levels (Figure S1G-K), confirming that organoids recovered from hypoxia.

A metabolic flux analysis was performed to functionally evaluate hypoxia’s effect on organoid glycolysis-TCA pathways (Figure 1H) by incubating organoids with ^13^C6-glucose in hypoxic or normoxic conditions for 3 hours. Mass spectrometry analysis of the glycolysis and TCA-cycle metabolites showed an increase in ^13^C labeling of glycolytic intermediates and a decrease in TCA cycle intermediates in hypoxic organoids. This suggests that in hypoxic organoids, glucose flux through glycolysis is increased while the flux of glycolytic intermediates through the TCA cycle is decreased. However, intracellular glucose levels remained relatively constant as evaluated by metabolic flux analysis and spatial metabolomics (Figure 1H-I). Spatial metabolomics also confirmed the hypoxia-induced shift from oxidative phosphorylation (decreased citrate, malate) to anaerobic metabolism (increased lactate) confirming that kidney organoid cells contain functional mitochondria which are affected by hypoxia. Taken together, these results indicate that hypoxic organoid model faithfully recapitulated acute and transient hypoxic conditions, providing an opportunity to examine kidney cell type-specific response.

### An organoid-based transcriptional signature of hypoxia identifies individuals with tubular injury and poor kidney disease outcome

We next evaluated the ability of our hypoxic kidney organoid model to capture kidney disease relevant gene expression. First, a 377-gene Hypoxic Organoid Signature was generated (Figure 2A) by applying stringent expression filters (Log_2_ fold change≥1, q-value≤0.01) to the differentially expressed (DE) genes from Figure 1F. When compared to a 155-gene Curated HIF gene set based on literature review and filtered by expression in kidney cell types (Table ST1), there was a 43-gene overlap. 144 of 155 Curated HIF Signature genes were expressed in organoids, with summary expression higher in hypoxic compared to normoxic kidney organoids (Figure 2B), providing evidence of model integrity reflecting HIF-related gene activity, though not all genes were included in the Hypoxic Organoid Signature due to the stringent filters employed.

**Figure 2.**
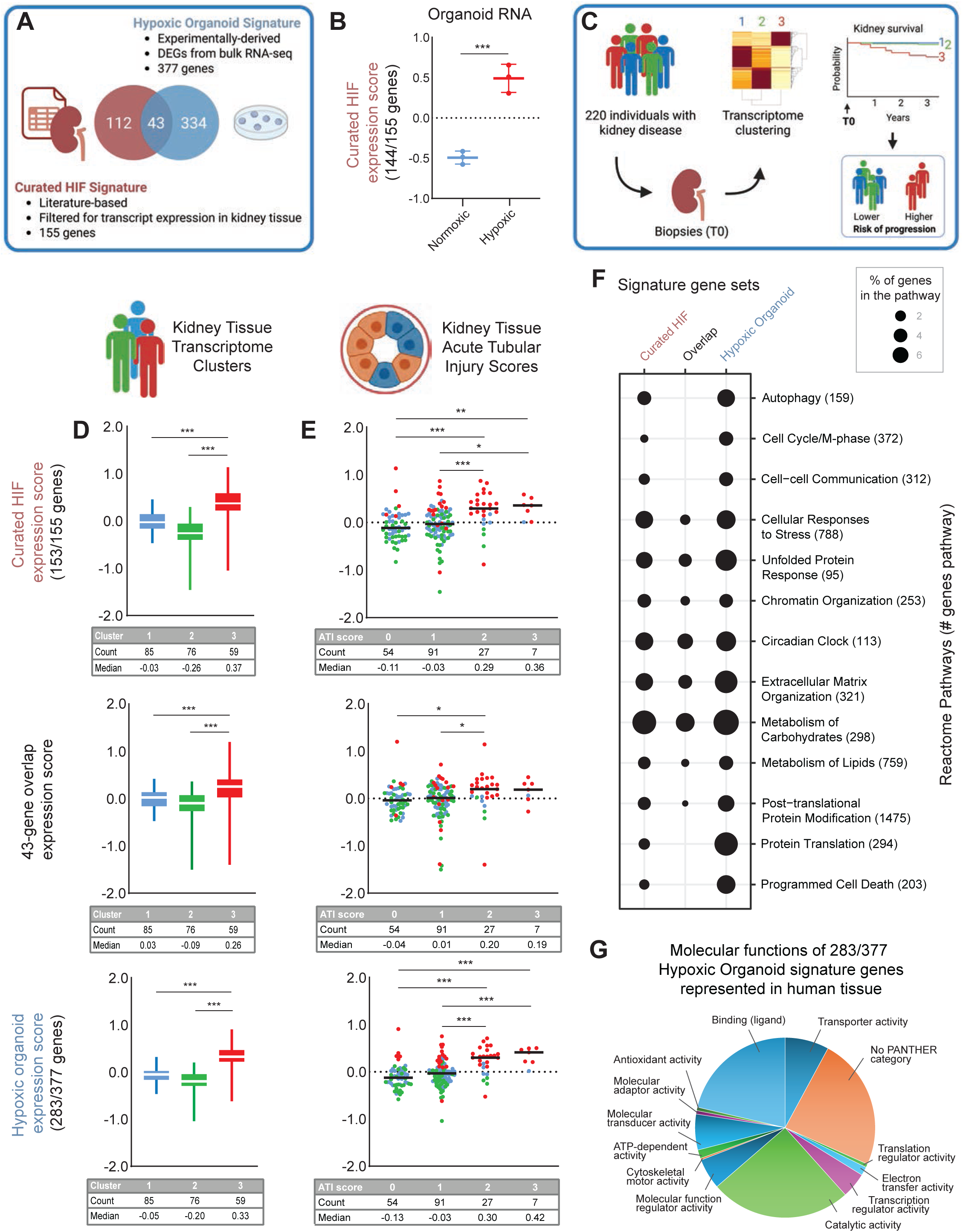
An organoid-based transcriptional signature of hypoxia identifies individuals with tubular injury and poor kidney disease outcome. **(A)** Venn diagram showing shared and unique genes of Hypoxic Organoid and Curated HIF Signatures. Created with BioRender. **(B)** Scatter plot of Curated HIF Signature summary gene expression in hypoxic and normoxic organoids. **(C)** Schematic of transcription-based clustering of tubulointerstitial kidney tissue from proteinuric individuals collected at time 0 (T0), with subsequent kidney survival by cluster. Created with BioRender. **(D)** Summary expression of gene signatures in (A) in kidney tissue transcriptome clusters in (C). Boxplots represent median (white line), interquartile range (colored box), and minimum and maximum values (whiskers). **(E)** Dot plot of individual tissue transcriptome summary gene expression (color corresponds to cluster in (D)), sorted by corresponding tissue acute tubular injury (ATI) score. Median expression, black line. Statistical comparisons between ATI scores shown; \**p*<0.05, \*\**p*<0.01, \*\*\**p*<0.001. **(F)** Bubble plot of gene contribution of Signature gene sets in (A) to Reactome pathways, represented as percent of total genes in pathway. **(G)** Pie chart of gene ontology (GO) Molecular Functions from the Panther Classification System represented by the 283 genes expressed in human kidney tissue (from D) of the 377-gene Hypoxic Organoid Signature. Data analyzed with unpaired t-test (B, D, E); \**p*<0.05, \*\**p*<0.01, \*\*\**p*<0.001. **Related data** Figure S2, Tables ST1-3.

Transcript levels of Signature genes were then assessed in micro-dissected tubulointerstitial kidney tissue obtained from a well-characterized cohort of individuals with proteinuric kidney disease (Figure 2C) to explore relevance to diseased kidney tissue. This previously described disease cohort self-sorted into three clusters of individuals based on transcript expression detected within the biopsied kidney tissue; cluster 3 (red) representing the group of individuals whose kidney disease was associated with the highest rate of progression within 40 months post-biopsy^25^. Summary gene expression scores for the Curated HIF, 43-gene overlap, and Organoid Hypoxia Signatures as well as both Signature’s unique genes were assessed in the clusters (Figures 2D, S2A-B). Each of these genes sets segregated cluster 3 from the other two tissue clusters indicating that HIF-related and hypoxia-related gene expression was higher in tissue of individuals whose disease progressed indicating that both the unique and overlapping components of the curated and experimentally derived gene sets captured disease relevant network activity.

Summary Signature gene set expression was then assessed relative to histologic evidence of tubular injury in kidney tissue from these same individuals for whom data were available (Figure 2E). Acute tubular injury (ATI) scores of kidney tissues were calculated based on the percentage of kidney cortex that demonstrated evidence of tubular injury (Figure S2C)^26,27^; a higher score correlated with more interstitial fibrosis, tubular atrophy and *HIF1A* transcript expression (Figures S2D-F) confirming that the ATI score correlated with tubular injury severity *in vivo*. While tubular injury scores were readily distinguishable by both Signatures’ collective gene expression scores, the 43-overlapping gene set was less effective, suggesting that each of the Signatures’ unique genes captured gene expression relevant to a spectrum of tubular injury (Figures S2G-H).

To determine what cell-based biological processes were captured by the Signature gene sets, they were explored using the Reactome Pathway Knowledgebase (Figures 2F, S2I, Tables ST2,4). In addition to Metabolism of Carbohydrates pathway, the Unfolded Protein Response, Programmed Cell Death, Protein Translation and Extracellular Matrix Organization pathways were prominently represented in the Hypoxic Organoid Signature and to a lesser extent in the Curated HIF Signature. Meanwhile, the 43-gene overlap did not capture most of these pathways indicating that each Signature adds genes uniquely to those pathways. The molecular functions represented by the Hypoxic Organoid Signature genes were also assessed for GO Molecular Functions (Figure 2G, Table ST3). Relevant to the Reactome pathways, this mechanism identified ATP-dependent, Electron transfer, Antioxidant, and Cytoskeletal motor activities as being represented. In the 94 genes of the Hypoxic Organoid Signature not significantly expressed in human kidney tissues (Figure S2J, Table ST5), Ligand binding, Catalytic and Transporter activity were predominant but also well represented in Figure 2G.

Together, these results indicate that the Hypoxic Organoid Signature captured gene activity that correlated with both poor long-term outcomes for individuals with proteinuric kidney disease, and histological evidence of significant tubular injury. Critically, as the Curated HIF Signature gene set also captured these elements, this supports the premise that the Hypoxic Organoid Signature gene set includes HIF-related gene activity. However, the Hypoxic Organoid Signature significantly expands the genes relevant to kidney disease and tubular injury providing additional information about kidney cell-based activities impacted by hypoxia.

### Hypoxia induces tubular epithelial cell apoptosis, alters cell energetics and enhances fibrosis

Based on correlation of the transcriptional activity of hypoxic kidney organoid with histological evidence of kidney tissue-level tubular injury, we reexamined the organoid tubules. Hypoxic organoids exhibited a significant increase in apoptotic tubular cells, decrease in phospho-S6, and increase in lipid droplet accumulation (Figures 3A-C, S3A) consistent with hypoxia-mediated organoid tubular cell injury and altered energetics. To explore whether hypoxic organoids developed fibrosis, we adapted the model in Figure 1A based on our prior results demonstrating that aging organoids develop fibrosis^28^; this required a hypoxic exposure earlier in culture to allow comparison by D25 (Figure 3D). Comparison of transcriptional profiles of the two models in Figures 1A and 3D at the end of 24 versus 48 hours of hypoxia confirmed significant overlap of gene expression seen in Figure 1F, as well as the expected fall in culture glucose level, and increase in *TGFB1* and *HAVCR1* expression (Figure 3E-F, S3B-C). However, tubular KIM-1 was unchanged (Figure S3D). Comparison of the bulk transcriptional profiles of the organoids immediately after 2 days of hypoxia versus 5 days of reoxygenation show that most mitochondrial and glycolytic pathway transcripts as well as the Hypoxic Organoid Signature score reverted to normoxic levels (Figures 3E-F, S3E). However, a few transcripts were significantly DE at D25 including inflammation-associated *ICAM1* and *PTX3* (Figure 3F, bottom).

**Figure 3.**
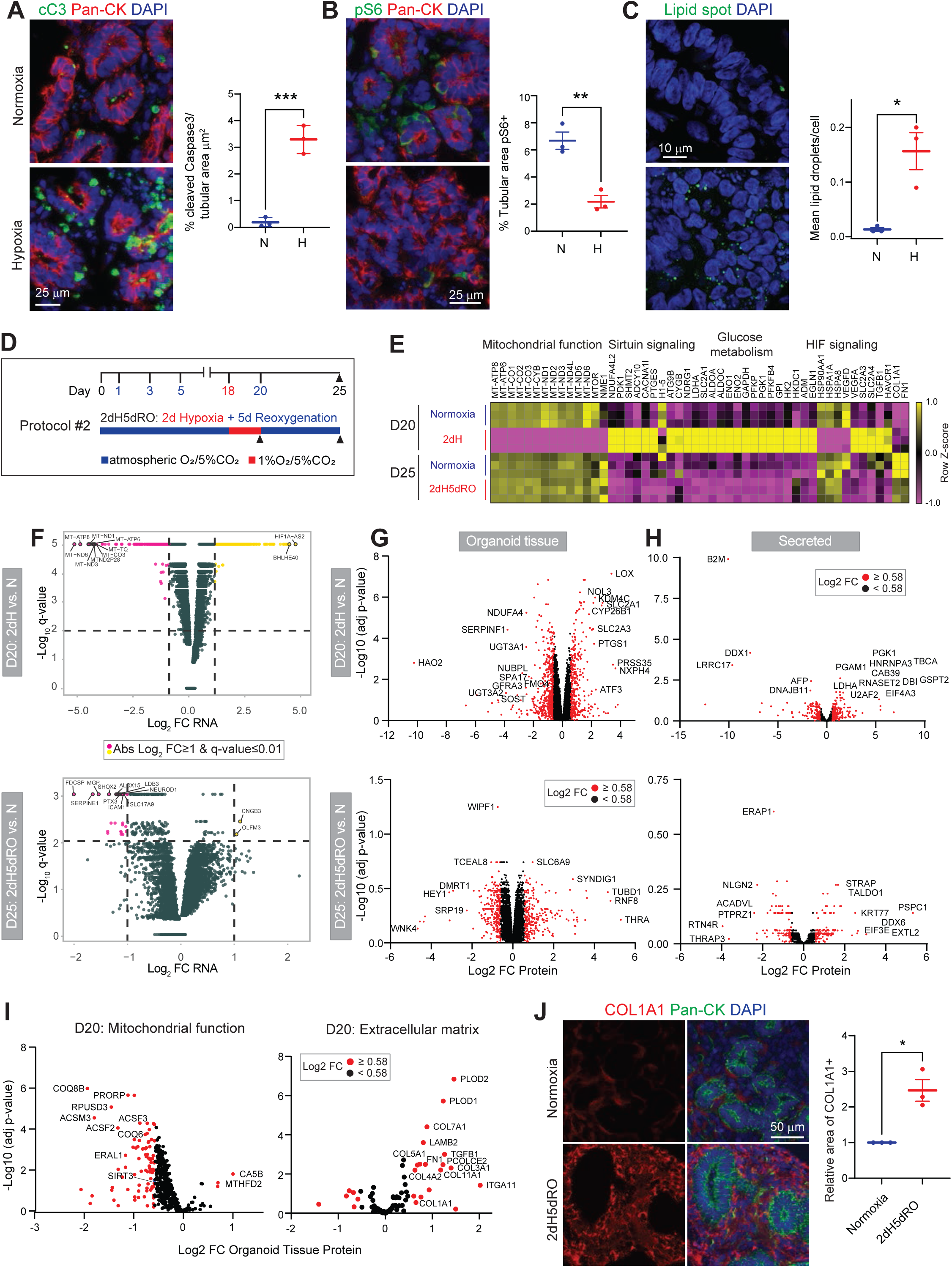
Hypoxia induces tubular epithelial cell apoptosis, alters cell energetics and enhances fibrosis. **(A-C)** Immunofluorescence and quantification (n=3) comparing expression of cleaved caspase 3 (A), phospho-S6 (B) and lipid accumulation (C), in normoxic and hypoxic organoids. (**D**) Schematic of Hypoxia Protocol #2 which includes 2 days of hypoxia followed by 5 days of reoxygenation. Black arrowheads indicate sample collection times at D20 (post-hypoxia) and D25 (post-reoxygenation). (**E**) Heatmap of DE genes (FDR<0.01) as in Fig. 1G in organoids treated with Hypoxia Protocol #2. (**F**) Volcano plots of DE genes in hypoxic vs. normoxic organoids at D20 acutely after hypoxia (top) or D25 after reoxygenation (bottom). DE transcripts meeting the significance threshold are represented by colored dots, with increased (yellow, upper right quadrant) or decreased (pink, upper left quadrant) relative expression; selected highly DE transcripts are labeled. Y axis maximum adjusted for visualization purposes. **(G and H)** Volcano plots of DE proteins in hypoxic vs. normoxic organoids treated with Hypoxia Protocol #2 as detected in tissue lysates (G) or (secreted into) culture media (H) at D20 (top) or D25 (bottom). Red dots represent proteins meeting the significance threshold. **(I)** Volcano plots of selected DE proteins at D20 analyzed in (G, top) with special focus on mitochondrial (left) and extracellular matrix (right) associated proteins. **(J)** Immunofluorescence and quantification (n=3) comparing COL1A1 expression in hypoxic following reoxygenation vs. normoxic organoids at D25. Data displayed as mean +/- SEM; n=biological replicates; Unpaired t-test (A, B, C), Welch’s t-test (J); \**p*<0.05, \*\**p*<0.01, \*\*\**p*<0.001. IF markers: nuclei (DAPI), tubular epithelial membrane (pan-CK); white scale bars, as labeled. **Related data** Figure S3.

Proteomics profiles of the organoid tissues and secreted proteins captured in parallel to the bulk transcriptomics samples revealed that the robust differential protein expression changes seen immediately after hypoxia also normalized after 5 days of reoxygenation, though many proteins still showed absolute Log_2_ fold changes in expression ≥0.56 (Figures 3G-H). A closer examination of the D20 tissue protein expression profile revealed a significant decrease in proteins associated with mitochondrial function and an increase in those associated with extracellular matrix (ECM) (Figure 3I). Five days after reoxygenation, hypoxic organoids showed significantly increased deposition of COL1A1 and trend in FN1 (Figures 3J, S3F) consistent with hypoxia-induced fibrosis.

These results confirm injury and altered energetics in tubular cells following hypoxia as well as persistence of gene expression alterations following reoxygenation. Notably, differential protein expression was detected for six transcripts DE at D25: CCN1, PPP1R17 were decreased in transcript and tissue protein; CNTN2, PTX3 and SERPINE1 were decreased in transcript and secreted protein; OLFM3 was increased in transcript and secreted protein (Figure S3G). Overall, these results are consistent with a decrease in matrix remodeling, apoptosis and senescence following hypoxia and reoxygenation which exceeds normoxic organoids, despite increased evidence of tissue fibrosis, which may reflect a hypoxic conditioning effect^29^.

### Hypoxia stimulates unfolded protein response and suppresses expression of mitochondrial β-oxidation proteins

As both unfolded protein response (UPR) and mitochondrial stress are integrally involved with apoptosis^30,31^, we examined hypoxia’s effect on genes involved in UPR and mitochondrial β-oxidation. Acutely hypoxic organoids significantly increased key proteins involved in UPR including ATF3 (Figure 4A). An outlier with a significant decrease was ATF6B which is thought to have an opposing role to the activating role of ATF6, a potent activator of endoplasmic reticulum (ER) stress response genes^32^. Hypoxia induced tubular cell expression of *XBP1* and *HERPUD1*, two additional key UPR genes, as shown by single cell transcriptional profiling (Figures 4C, S4A-C).

**Figure 4.**
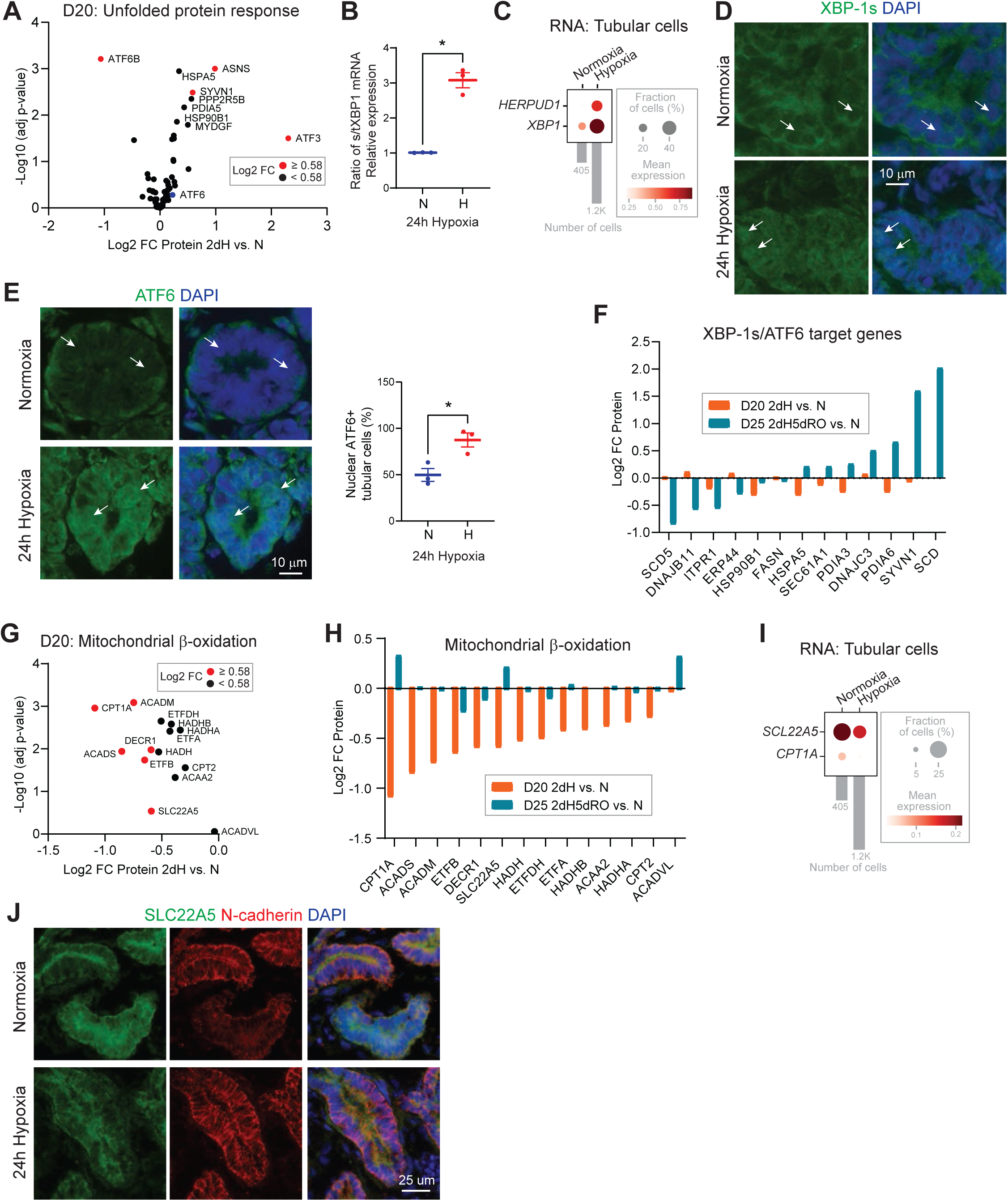
Hypoxia stimulates unfolded protein response and suppresses expression of mitochondrial β-oxidation proteins. **(A)** Volcano plot of DE proteins of the UPR pathway in D20 hypoxic vs. normoxic organoids. Red dots represent proteins meeting the significance threshold. **(B)** Graph comparing spliced (s) isoform to total (t) *XPB1* expression levels in acutely hypoxic and normoxic organoids (n=3). **(C)** Dot plot of transcriptional expression of key UPR-related genes in 24h hypoxic and normoxic tubular cells from single cell profiled organoids. **(D and E)** Immunofluorescence comparing nuclear localization of XBP-1s (D) and ATF6 with quantification (E, n=3) in hypoxic organoid tubular cells (based on nuclear pattern). White arrows highlight nuclei (DAPI) for comparison between conditions. **(F)** Bar plot comparing differential expression of XBP-1s and ATF6 target proteins in hypoxic vs. normoxic organoids at D20 post-hypoxia (orange bars) and D25 post-reoxygenation (blue bars). **(G)** Volcano plot of DE proteins related to mitochondrial β-oxidation, comparing D20 acutely hypoxic to normoxic organoids. Red dots represent proteins meeting the significance threshold. **(H)** Bar plot comparing differential expression of mitochondrial β-oxidation proteins in hypoxic vs. normoxic organoids at D20 post-hypoxia (orange bars) and D25 post-reoxygenation (blue bars). **(I)** Dot plot of transcriptional expression of *SLC22A5* and *CPT1A* in 24h hypoxic and normoxic tubular cells from single cell profiled organoids. **(J)** Immunofluorescence comparing SLC22A5 expression in hypoxic and normoxic organoids. Data displayed as mean +/- SEM; n=biological replicates; Welch’s t-test (B), unpaired t-test (E); \**p*<0.05. IF markers: nuclei (DAPI), tubular epithelial membrane (N-cadherin); white scale bars, as indicated. **Related data** Figure S4.

Functional evidence of UPR pathway activation in hypoxic organoids included increased level of spliced isoform of *XBP1* (*sXBP1*) (Figure 4B, S4D) and nuclear localization of XBP1-s and ATF6 in tubular cells (Figures 4D-E). While protein levels of several XBP1-s and ATF6 target genes were unaffected by acute hypoxia, there was more variability after 5 days of reoxygenation despite a decrease in *sXBP1* level and nuclear ATF6 localization (Figures 4F, S4E-F).

Notably, as opposed to UPR proteins, mitochondrial proteins involved in β-oxidation were potently decreased with acute hypoxia but generally normalized following reoxygenation (Figures 4G-H). Two proteins involved in the carnitine shuttle, CPT1A and SLC22A5, as well as their transcript levels, and protein level of SLC22A5 were also decreased in hypoxic tubular cells (Figures 4H-J), suggesting a mechanism involved in altered lipid metabolism resulting in the hypoxia-induced accumulation of lipid droplets observed in Figure 3C. Thus, the hypoxic organoid-derived protein data align with the human kidney-based transcriptional data, confirming the relevance of the hypoxic organoid model to tubular injury in proteinuric kidney disease shown in Figure 2.

### Dapagliflozin protects kidney tubular epithelial cells from hypoxia-mediated injury via decrease in reactive oxygen species

Next, we employed the highly selective SGLT2 inhibitor dapagliflozin^33^ to test the hypothesis that SGLT2 inhibition protects tubular cells in hypoxic conditions. Given the low level of SGLT2 expression in organoid tubular cells (Figure 5A), we anticipated the experimental impact of SGLT2 inhibition would be minimal. However, the addition of dapagliflozin to hypoxic organoids resulted in a marked increase in organoid culture media glucose levels, while organoid tissue levels of glucose decreased (Figure 5B-C). Meanwhile high levels of hypoxia-induced *SLC2A1* and GLUT-1 expression remained unchanged in tubular cells, as did expression of another glucose transporter *SLC2A3* (Figure S5A-B). An alternative explanation for the observed dapagliflozin-induced change in tissue and media glucose levels is a suppressive effect on cell energetics. To this point, dapagliflozin further suppressed hypoxia-mediated phospho-S6 levels, consistent with decreased mTOR pathway activity, and further increased accumulation of lipid droplets, consistent with altered lipid metabolism (Figure 5D-E).

**Figure 5.**
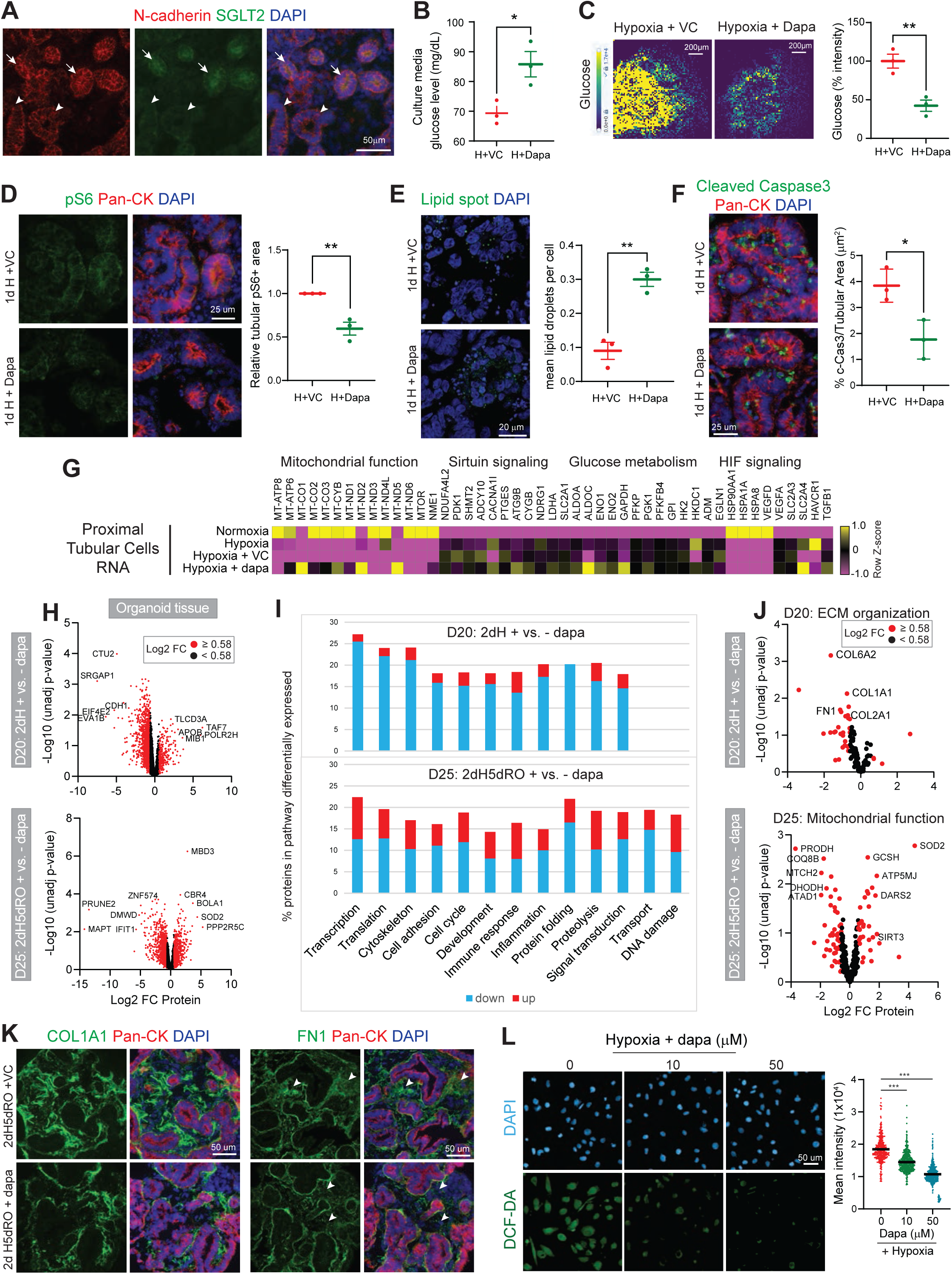
Dapagliflozin protects kidney tubular epithelial cells from hypoxia-mediated injury via decrease in reactive oxygen species. **(A)** Immunofluorescence showing SGLT2 expression detected in normoxic organoid tubular cells. **(B)** Graph comparing culture media glucose levels of organoids after 24h hypoxia +/- dapagliflozin (n=3). **(C)** MALDI-MSI and quantification (3 organoids) of glucose analyte in organoids after 24h hypoxia +/- dapagliflozin. Yellow, highest signal intensity. **(D-F)** Immunofluorescence comparing detection of phospho-S6 (D), lipid accumulation (E), cleaved caspase 3 levels (F) with corresponding quantification graphs (n=3), in organoids after 24h hypoxia +/- dapagliflozin. (**G**) Heatmap comparing transcriptional expression of selected genes in proximal tubular cells as in Fig. 1G in 24h hypoxic +/- dapagliflozin and normoxic single cell profiled organoids. (**H**) Volcano plots of DE proteins in tissue of D20 post-hypoxic (top) or D25 post-reoxygenated (bottom) organoids +/- dapagliflozin. (**I**) Stacked bar plots comparing dapagliflozin’s effect on predicted pathway activity based on differential protein expression in D20 post-hypoxic and D25 post-reoxygenation organoids, as assessed by Metacore Process Networks (FDR<0.05). (**J**) Volcano plots of DE proteins detected in (H) highlighting differential expression of ECM (top) and mitochondrial (bottom) proteins, at D20 and D25 respectively. (**K**) Immunofluorescence comparing dapagliflozin’s effect on COL1A1 and FN1 expression in hypoxic-then-reoxygenated D25 organoids as in (H). (**L**) Immunofluorescence comparing dapagliflozin’s effect on ROS generation (green) in hypoxic HK2 cells treated for 24h, with quantification (n=1 +/- SD, technical triplicates). Data displayed as mean +/- SEM; n=biological replicates; Unpaired t-test (B, C, E, F), Welch’s t test (D), One-way Anova (L); \**p*<0.05, \*\**p*<0.01, \*\**p*<0.001. IF markers: nuclei (DAPI), tubular epithelial membrane (N-cadherin/pan-CK); white scale bars, as indicated. **Related data** Figure S5, Table ST6.

Notably, dapagliflozin also significantly reduced hypoxia-induced tubular cell apoptosis (Figure 5F). To further explore mechanisms that contributed to this beneficial result, single cell transcriptional profiling of organoid PT cells was performed and showed that dapagliflozin reversed the hypoxia-induced reduction in mitochondrial gene expression, with expression approaching normoxic levels (Figures 5G, S4A). However, nuclear localization of HIF-1, transcript levels of genes involved in HIF-1 and sirtuin signaling, glucose metabolism as well as *TGFB1* and *HAVCR1* were all unchanged, as were levels of *sXBP1* and nuclear localization of ATF6 (Figures 5G, S5C-F). These results indicate that dapagliflozin did not significantly affect hypoxia-induced HIF-1- and UPR-related transcriptional programs.

Given the limited dapagliflozin-induced transcriptional changes for those genes assessed, we next explored its effect on protein levels in hypoxic organoids (Figures 5H, S5I). Although differential expression of proteins between dapagliflozin- and control-treated hypoxic organoids did not achieve adjusted p-values of ≤ 0.05 as seen with hypoxia versus normoxia samples (Figure 3G-H), inclusion of samples assessed in triplicate in 2 independent experiments supported our use of relative protein levels with unadjusted p-values ≤ 0.05. Dapagliflozin treatment resulted in an overall significant shift towards decreased protein expression in acutely hypoxic organoids (compare Figure 5H to 3G). Consistent with these results, process networks represented by these proteins (Figure 5I (top), Table ST6) involved in transcription, translation and protein folding were markedly decreased. After 5 days of reoxygenation, dapagliflozin still markedly affected differential expression of proteins, but with a more balanced profile of network activity; networks involved in cytoskeleton, cell adhesion, signal transduction and cell cycle activities all increased, consistent with dapagliflozin-enhanced cell recovery.

Closer examination of proteins involved in ECM organization (Figure 5J) revealed that expression of COL1A1 and FN1 decreased with dapagliflozin treatment despite hypoxic conditions (Figures 5J-K), consistent with an anti-fibrotic effect. After reoxygenation, expression of two key proteins involved in mitochondrial stress response, SIRT3 and SOD2, increased with dapagliflozin. A protective role for dapagliflozin against ROS generation under hypoxic conditions was confirmed in HK2 proximal tubular cells (Figure 5L). Meanwhile, expression of proteins involved in glucose metabolism was modestly decreased by dapagliflozin with acute hypoxia which persisted 5 days following reoxygenation suggesting persistent alteration of cell energetics (Figure S5G, compare orange to blue bars). UPR protein levels were minimally affected (Figure S5H), in line with the observed transcriptional response.

Taken together, these results indicate that dapagliflozin imparts a significant beneficial effect in hypoxic tubular epithelial cells related to improved resiliency to mitochondrial stress, mediated through enhanced mitochondrial gene expression and functional response to ROS. Without the compounding negative effect of ROS generation, an active UPR program increases tubular cell survival in hypoxia.

### Integrated transcriptomic and proteomic analysis reveals markers of dapagliflozin-mediated modulation of hypoxia-induced cellular stress response

We posited that dapagliflozin would reverse the expression of genes from the 377-gene Hypoxic Organoid Signature in Figure 2, and identify additional cellular functions affected by dapagliflozin. Furthermore, the ability to align experimental data with available transcriptional human kidney data would enhance modeling and treatment goals. However, the Hypoxic Organoid Signature gene expression was not significantly affected by dapagliflozin, in either acute hypoxia or reoxygenation conditions or in either proximal tubular cells or whole organoids (Figure S6A-C).

Given these results, we adjusted our strategy to leverage the dapagliflozin-treated hypoxic organoid data in Figure 5 to identify highly DE proteins within the 377-gene set with which we could reapproach human transcriptional data and evaluate for correlation with organoid expression. As a first step, organoid protein levels of genes that were transcriptionally DE in individuals with T2D when treated with SGLT2-i^34^ were assessed. With acute hypoxia, the pattern of expression for glycolysis and TCA cycle pathway proteins in dapagliflozin-treated organoids reversed, consistent with what was observed in the SGLT2-i treated T2D individuals (Figure 6A). However, after reoxygenation, this difference between treatment and control organoid protein levels disappeared suggesting that the acute hypoxia organoid condition more closely reflected *in vivo* physiology in this group. These results reinforced the pursuit of targeted assessment of gene expression within the Hypoxic Organoid Signature.

**Figure 6.**
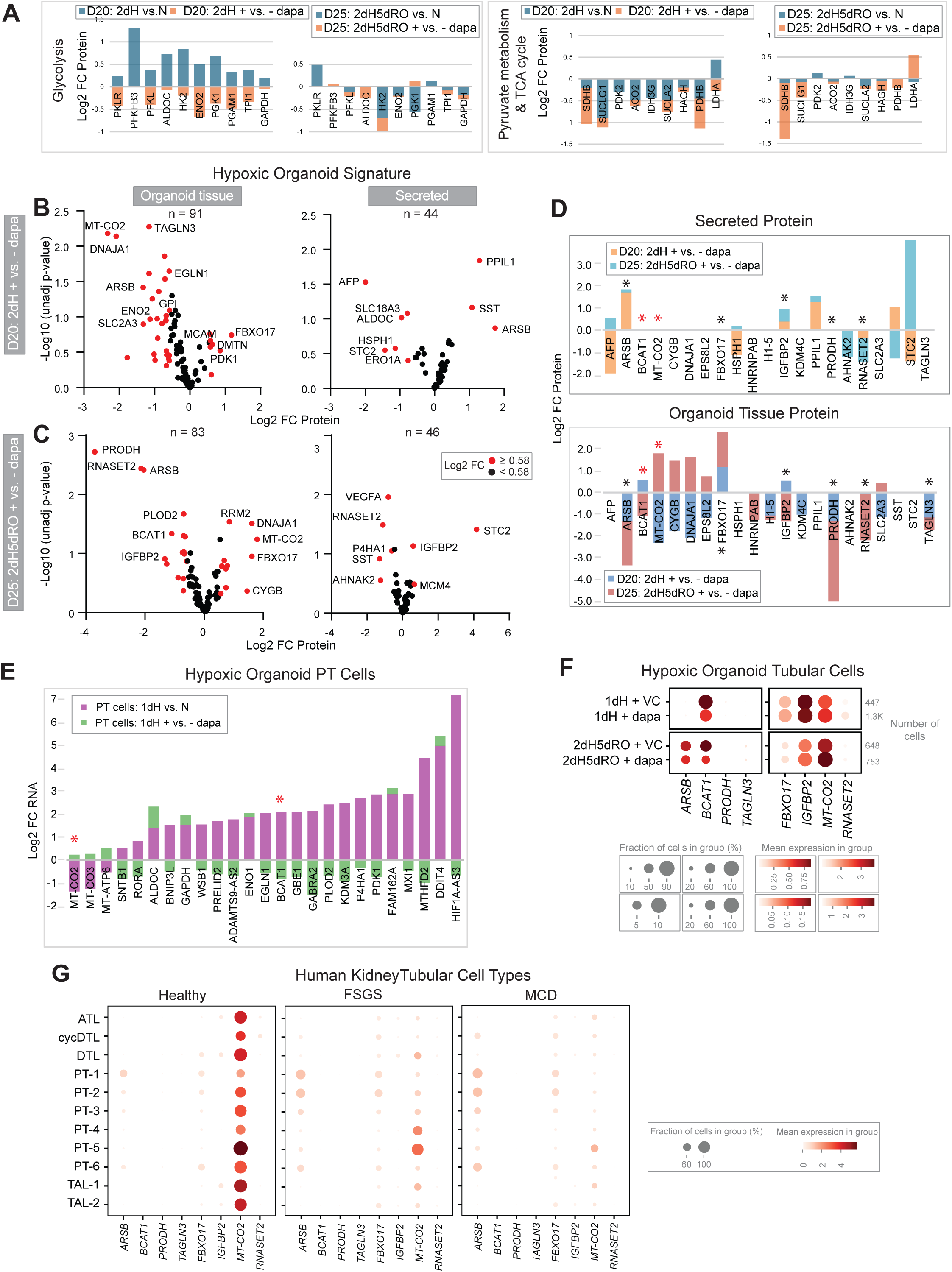
Integrated transcriptomic and proteomic analysis reveals markers of dapagliflozin-mediated modulation of hypoxia-induced cellular stress response. **(A)** Stacked bar plots comparing the effect of dapagliflozin on differential expression of glycolysis, pyruvate metabolism and TCA cycle-related proteins in D20 post-hypoxic (left graphs) and D25 post-reoxygenated (right graphs) organoids (orange bars), to the effect of hypoxia on differential expression of proteins relative to normoxia (blue bars) at the same time points. **(B and C)** Volcano plots comparing dapagliflozin’s effect on hypoxic organoid tissue or secreted differential protein expression of 377 Hypoxic Organoid Signature genes at D20 post-hypoxia (B) and D25 post-reoxygenation (C) +/- dapagliflozin. **(D)** Stacked bar plots comparing the top 21 DE proteins in secreted (top) and organoid tissue (bottom) compartments, at D20 post-hypoxia and D25 post-reoxygenation, +/- dapagliflozin. Asterisks, DE proteins with Log_2_ FC ≥ 1 in at least 1 timepoint or compartment (secreted or tissue-based). **(E)** Stacked bar plots showing the effect of dapagliflozin on top 25 hypoxia-induced DE genes in organoid proximal tubular cells; compare 24h hypoxic +/- dapagliflozin (green bars) to 24h hypoxic vs. normoxic organoids (purple bars). Red asterisks, genes with protein (D) and transcript expression affected by dapagliflozin. **(F)** Dot plots comparing dapagliflozin’s effect on hypoxic organoid tubular cell transcriptional expression of genes of interest from (D & E); post-hypoxia and post-reoxygenation. **(G)** Dot plots comparing transcriptional expression of genes of interest (F) in tubular cell types in individuals with FSGS and MCD (as in Figure 2) relative to healthy kidneys. **Related data** Figure S6.

Protein expression in hypoxic organoids exposed to dapagliflozin was detected for a fraction of Signature genes (n=102 tissue and secreted), while a smaller fraction was DE (n=61) (Figure 6B-C). However, examination of the transcripts encoding this smaller subset of DE proteins had minimal effect on the dapagliflozin-treated Hypoxic Organoid Signature score (Figure S6D). Graphing the differential expression levels of protein versus bulk RNA in hypoxic dapagliflozin-versus control-treated organoids (for the genes expressed in both) revealed significant discordant expression changes (Figure S6E). These results highlight the challenges of aligning transcriptional and protein effects in this setting, as well as the benefit of our integrated approach to assess dapagliflozin’s impact on cellular functions.

We next narrowed our focus to the most highly DE genes. Dapagliflozin-induced DE proteins with absolute Log_2_ FC≥1 (n=21) in either tissue or secreted proteome of either acutely hypoxic or reoxygenated organoids were compared to the top 25 most DE transcripts in hypoxic PT cells (Figure 6D-E). Two genes (red asterisks) with organoid tissue protein levels reversed by dapagliflozin following reoxygenation, BCAT1 and MT-CO2, were detected as DE transcripts in dapagliflozin-treated hypoxic organoid PT cells. Six other highly DE proteins were also selected (Figure 6D, black asterisks) for further investigation: 3 were present in both organoid tissue and secretome (RNASET2, IGFBP2 and ARSB), while 3 were expressed in organoid tissue only (PRODH, TAGLN3 and FBXO17).

The transcriptional expression pattern of these eight genes showed that six were expressed in kidney organoid tubular and human kidney tubular cell subtypes in individuals with proteinuric kidney disease from the same cohort^35^ as in Figure 2 (Figures 6F-G, S6G-H); PRODH and TAGLN3 were nominally expressed in tubular cells of either dataset. Note that proximal and distal tubular cell types were included in this transcriptional comparison to both increase the number of transcripts (for organoids) and assess tubular expression pattern (for human tissue).

This integrated proteome-transcriptome analysis of dapagliflozin-treated hypoxic organoids identified six genes of the original 377-gene Hypoxic Organoid Signature set that were potently differentially expressed in both hypoxic kidney organoid and diseased human kidney tubular cell types and were experimentally reversed by dapagliflozin. FBXO17 was the only gene included in the 155-gene Curated HIF Signature (Figures 2A, S6F). A unifying theme for these affected genes is metabolic activity with dapagliflozin resulting in: decreased ARSB (breaks down glycosaminoglycans, silencing mimics hypoxia)^36,37^ and RNASET2 (endoribonuclease involved in degradation and processing of mitochondrial and non-coding RNA)^38,39^; increased FBXO17 (recognizes and binds denatured glycoproteins in an E3-ubiquitin ligase complex)^40,41^; acutely increased then decreased BCAT1 (catalyzes transamination of branched-chain amino acids, increased expression driven by hypoxia and HIF activity)^42,43^ and IGFBP2 (binds and regulates insulin-like growth factor, proapoptotic, expression increased by hyperglycemia)^44,45^; and acutely decreased then increased expression MT-CO2 (mediates transfer of electrons from cytochrome C to oxygen, hypoglycemia increases expression)^46,47^. While the directional effect of some of these genes’ transcript and protein levels are consistent between acute hypoxia and reoxygenation, the reversal of expression profile of others suggests an opportunity to identify unique profiles reflecting acute versus subacute hypoxic cell injury.

## DISCUSSION

To investigate mechanisms of hypoxic kidney cell injury, we generated a robust model using hPSC–derived kidney organoids and rigorously validated the response to hypoxic conditions by confirming HIF-1 associated gene expression and showing altered glucose metabolism and TCA cycle function. Importantly, we showed that organoids functionally recovered following hypoxic exposure. A set of highly differentially expressed transcripts, many not previously described in hypoxia-mediated kidney injury, segregated a sub-group of individuals with proteinuric kidney disease at highest risk for progression and correlated with histological evidence of tubular injury in kidney tissue, demonstrating that this model captured key cellular processes involved in clinically relevant kidney cell injury. Moreover, these results confirm that hypoxia-related gene expression is an integral component of proteinuric kidney disease at risk for disease progression. Analysis of the hypoxic organoids revealed enhanced tubular cell apoptosis associated with UPR activity and ECM production and deposition and as well as altered energetics, mitochondrial gene expression and lipid metabolism.

Low SGLT2 expression and lack of vascular flow and glomerular filtration provided an opportunity to focus on dapagliflozin’s off target effects in kidney tubular cells. Dapagliflozin significantly attenuated hypoxia-induced tubular apoptosis by enhancing mitochondrial functionality; protein levels of SOD2 and SIRT3 increased which countered ROS generation. Additionally, hypoxia-induced ECM production and deposition decreased with dapagliflozin. Conversely, dapagliflozin’s effect on expression of hypoxia-induced transcriptional profile was relatively modest with HIF-1α localization persisting in the nucleus, suggesting a more nuanced effect on hypoxia-induced HIF-1-associated transcriptional activity relative to a more robust effect on post-transcriptional activity. An integrated proteomics-transcriptional analysis, however, identified six hypoxia-altered proteins associated with cell metabolic activity whose expression was significantly impacted by dapagliflozin. Transcriptional analysis confirmed expression of this gene set in tubular cells of organoids and reversed by dapagliflozin, while expression of this gene set in tubular cells of human kidneys was altered in disease. Together, these results identified a robust kidney cell response to hypoxia and provided integrated experimental evidence of beneficial effect of dapagliflozin on kidney tubular cell survival following hypoxic exposure.

We did not observe a dapagliflozin effect on hypoxia-induced nuclear localization or overall cell level expression of HIF-1α which contrasts with the results reported by Bessho et al.^48^; in that study, hypoxia-mediated HIF-1α levels were reversed by luseogliflozin treatment of human renal proximal tubular epithelial cells (HRPTECs) and proximal tubules in type 2 diabetes model *db/db* mice. This may be related to reported variable effects among SGLT2-i compounds^49,50^. Indeed, luseogliflozin did not decrease albuminuria in *db/db* mice^51^ as was observed with dapagliflozin^52^. Additionally, we did not observe a dapagliflozin effect on hypoxia-induced UPR protein expression; instead, we observed enhanced expression of SOD2 and SIRT3, as well as suppressed ROS generation. Our results contrast with those reported by Shibusawa et al.^53^ in which dapagliflozin protected against C2 ceramide induced ER-stress mediated apoptosis in HK2 cells & *db/db* mice; notably, a potentiating role of mitochondrial stress response was not investigated in these studies. However, our results are consistent with a protective effect of empagliflozin on hypoxia-mediated suppression of SIRT3 expression; loss of SIRT3 contributed to aberrant glycolysis, HIF-1α stabilization, increased ROS generation and epithelial to mesenchymal transition^54–56^.

Targeted examination of urinary biomarkers in humans to better define SGLT2-i’s beneficial effects on kidney physiology^57^ provided limited insight in part because causality could not be established. Our use of a simplified model system of kidney cells provided the ability to discern the experimental effect of dapagliflozin on tubular cell types. Our organoid-based results can be integrated with human studies to support development of biomarkers of dapagliflozin’s effects. For example, one of the genes that we identified in Figure 6 as impacted by dapagliflozin in BCAT1 which catalyzes the first step of branched chain amino acid catabolism^58^; dapagliflozin has been shown to enhance urinary excretion of BCAA-derived metabolites in humans and mice without changes in plasma levels, consistent with enhanced kidney BCAA-catabolism^59^.

A significant knowledge base of gene activity relevant to hypoxia is available, information that was critical in developing our hypoxia kidney model. However, it largely stems from cancer research^60^ and does not adequately capture gene expression relevant to hypoxia-mediated kidney cell injury. Our identification of a significantly expanded set of genes relevant to acute hypoxia-mediated kidney cell injury and confirmed relevance of our organoid model to proteinuric kidney disease. Interestingly, acute hypoxia gene activity of organoids was able to segregate a subgroup of proteinuric individuals with progressive kidney disease, suggesting that acute hypoxia is an active process in this chronic disease. Whether this gene expression pattern also reflects gene activity of chronic hypoxia or are a harbinger of the transition for AKI to CKD is to be determined.

Current clinical diagnosis of AKI relies on rise in serum levels of creatinine or decrease in urine output. Our identification of hypoxia-mediated kidney cell injury related gene activity could enhance biomarker development leading to improved identification of individuals with kidney injury and assessment of severity. Further, we identify potential gene candidates to gauge clinical therapeutic response. Our results suggest a rationale for continuing dapagliflozin during potential acute kidney hypoperfusion- and hypoxia-inducing clinical scenarios but will require clinical investigation, especially as to whether it is a protective or therapeutic intervention or reflects effects of other SGLT2-i compounds.

This robust, rigorously characterized model of hypoxia-mediated kidney cell injury and associated data will be available for use by the research community. Collectively, this model represents a valuable platform for investigating pathogenic mechanisms of kidney disease and identifying novel therapeutic targets. Moreover, cross species comparison can be performed to identify which non-human model systems reflect gene activity relevant to human kidney disease.^61^

### Limitations and Future Directions

The current paucity of dapagliflozin-treated human kidney disease-related data limits the ability to align organoid modeling with *in vivo* treatment effect at this point; however, we anticipate expansion of relevant datasets such that this will become feasible soon. In the meantime, we can use our current organoid-derived data to investigate gene activity in kidney cell types in available datasets of human AKI and CKD. An additional limitation is our use of a single SGLT-i, dapagliflozin, for this project. However, this limitation is off set by the depth and breadth of experimental data types that we generated for dapagliflozin, which can be used to compare with human kidney tissue datasets given dapagliflozin’s wide clinical use, and as a benchmark to investigate cellular effects of other SGLT2-i compounds, as well as additional therapies. Additionally, though our organoid model system ignores the impact of immune cells and hemodynamic and nephron-based physiology of the kidney, this approach provided the ability to discern kidney cell type specific effects of hypoxia. Future iterations of the organoid model can build on this knowledge base.

## SUPPLEMENTARY INFORMATION

Supplementary Figures S1-S6 and Legends

Supplementary Tables ST1-ST6

## ACKNOWLEDGEMENTS

Financial support was provided as follows: NIH funding UH3TR003288 (JLH, MK), K08DK124449 (JS) and R03DK138962 (JS), F30DK121463 (JB); Novo Nordisk Foundation funding NNF19OC0056043 and NNF24OC0095902 (MMR); Breakthrough T1D grant #5-COE-2019-861-S-B and gift funding (MK, SP, JLH). The authors wish to thank Breakthrough T1D and donors for their generous support of this research.

This project was supported by the University of Michigan Human Stem Cell Gene Editing Core Facility (RRID:SCR_026755), School of Medicine Advanced Genomics Core Facility (RRID:SCR_025788) and BRCF Microscopy Core Facility (RRID:SCR_026722), the O’Brien Kidney Centers (RRID:SCR_015270) and University of Michigan Kidney Translational Core Center (RRID:SCR_015903), the last two funded by NIH U54DK137314.

The Nephrotic Syndrome Study Network (NEPTUNE) is part of the Rare Diseases Clinical Research Network (RDCRN), which is funded by the National Institutes of Health (NIH) and led by the National Center for Advancing Translational Sciences (NCATS) through its Division of Rare Diseases Research Innovation (DRDRI). NEPTUNE is funded under grant number U54DK083912 as a collaboration between NCATS and the National Institute of Diabetes and Digestive and Kidney Diseases (NIDDK). Additional funding and/or programmatic support is provided by the University of Michigan, NephCure Kidney International, Alport Syndrome Foundation, and the Halpin Foundation. RDCRN consortia are supported by the RDCRN Data Management and Coordinating Center (DMCC), funded by NCATS and the National Institute of Neurological Disorders and Stroke (NINDS) under U2CTR002818.

## DATA AVAILABILITY

NEPTUNE bulk RNA-seq tubulointerstitium kidney biopsy data from NEPTUNE participants with FSGS/MCD were accessed via GEO accession number GSE182380. NEPTUNE OCEAN processed data available for visualization at https://neptune-ocean.cxg.miktmc.org/.

The following data are deposited in the NCBI Gene Expression Omnibus:

- sc RNA-seq:

- GSE306041 (D24_1d Hypoxia vs. Normoxia & Hypoxia+vs.-Dapa, D25_2dH5dRO+ vs. - Dapa)

BulkRNA-seq and Mass spectrometry proteomics data will be available after peer-review and publication.

## EXPERIMENTAL MODEL AND STUDY PARTICIPANT DETAILS

### Human Subjects

Publicly available data from the Nephrotic Syndrome Study Network (NEPTUNE, NCT01209000, HUM00158219) was used in this study. NEPTUNE is an existing observational study^62^ of individuals with proteinuric kidney disease from which kidney tissue for biopsies was micro-dissected into glomerular and tubulointerstitial compartments. Bulk tubulointerstitial transcriptional data from 220 NEPTUNE participants with FSGS/MCD^25^ were used in this study. Single cell and nuclear transcriptional profiling data from NEPTUNE participant and living donor kidney biopsy tissue^35^ was accessed on the Cell x Gene OCEAN platform (GEO data release embargoed pending publication), site software was used to generate visualization of data^63^.

### Stem cell lines

Studies with hPSCs were performed in accordance with University of Michigan’s Human Pluripotent Stem Cell Research Oversight Committee and NIH regulations. UM77-2 human (female) embryonic stem cell line (hESCs, NIHhESC-14-0278) was previously obtained through MStem Cell Laboratories. Breakthrough T1D-derived induced pluripotent stem cells (iPSCs) lines 49E (male) and 51A (female) (HUM00171983; HUM00185622) were reprogrammed from peripheral blood mononuclear cells isolated from donors with T1D at the University of Michigan’s Human Stem Cell & Gene Editing Core. Normal karyotype and expected sex for UM77-2s were confirmed by CNV assessment by Illumina bead array at University of Michigan’s Advanced Genomic Core. For T1D iPSCs, normal karyotyping and expected sex were confirmed by WiCell Research Institute.

## RESOURCES

Resources will be available after peer-review and publication.

## METHODS

### Kidney organoid generation

Kidney organoids derived from UM77-2 human embryonic stem cells (hESC) were generated according to previously described protocols^64,65^. Briefly, on D0, UM77-2 hESCs were dissociated with Accutase and seeded at a density of 15,000 cells per well in 24-well plate, precoated with 1% Geltrex in DMEM/F12, containing mTeSR1 and Y-27632 (10 µM). Media was replaced with 1.5% Geltrex in mTeSR1 on D1, mTeSR1 on D2, advanced RPMI supplemented with GlutaMAX (1x), CHIR99021 (10 µM) and Noggin (10 ng/mL) on D3. Cells were maintained for 36h, followed by a replacement with RB media [advanced RPMI supplemented with B27 supplement (1x), GlutaMAX (1x) and Penicillin-Streptomycin] on D5. Thereafter, RB media was replaced every third day until collection day. The same protocol was applied on T1D iPSCs with the exception that CHIR99021 was used at 12 µM for kidney organoid differentiation.

### Hypoxia and compound treatment

Two different protocols were applied for hypoxia treatment depending on the specific questions being asked (Figures 1A and 3D). Kidney organoids were placed in a hypoxia incubator chamber, with an oxygen monitor (OXY-2) and a petri-dish containing autoclaved water for adequate humidification (Figure 1A). The lid of hypoxia chamber was sealed with Parafilm and buckled with a ring clamp. The hypoxia chamber was purged with hypoxic gas consisting of 1%O_2_, 5%CO_2_ balanced N2 at a rate of 20 liters per minute for 4 min using a single flow meter and then placed in a cell culture incubator for the hypoxia treatment time. The oxygen level in the hypoxia incubator chamber was maintained at 1% throughout the experiment by re-purging the hypoxia chamber as needed (Figure S1A). In Hypoxia Protocol #1 (Figure 1A), kidney organoids at D23 were placed in the hypoxia chamber for 24h, and then samples were collected on D24. In Hypoxia Protocol #2 (Figure 3D), kidney organoids at D18 were placed in the hypoxia chamber for 48h and then were placed under normoxia for 5 days where samples were collected on D20 and D25. For the 48h hypoxia treatment, plates were double fed with media at the start of the experiment. Kidney organoids were treated with dapagliflozin (10 µM) in RB media at D23 for Protocol #1 and D18, D20 and D22 for Protocol #2. Vehicle control (VC) wells were treated with corresponding volume of DMSO, not to exceed 0.1%. At specified time points, samples were collected for subsequent processing by different techniques. Glucose levels in the kidney organoid supernatants were recorded with the GM100 Glucose Meter using Key 085 for RPMI media. In brief, 50 µl of supernatants were collected and were allowed to reach ambient temperature before measuring 2 readings/condition/replicate (20 µl/reading) from 3 independent experiments.

### Quantitative real-time PCR

Total RNA was collected from kidney organoid spheroids by scraping in ice-cold PBS, lysing in 500 µl of TRIzol, and extracting using Direct-zol RNA Miniprep kit. One µg of total RNA was reverse-transcribed into cDNA using SuperScript First-Strand Synthesis System for RT-PCR, as per manufacturer’s protocol. Quantitative real-time PCR (qRT-PCR) was performed in a QuantStudio 7 Flex Real-Time PCR System using TaqMan Fast Universal PCR Master Mix and TaqMan probes in a final volume of 10 µl per reaction. The ΔΔCq method was applied to calculate the relative expression of target gene after normalization to the housekeeping gene, 18S. For spliced *XBP1* (*sXBP1*), normalization was performed using total *XBP1* (*tXBP1*) levels.

### Immunofluorescence and quantification

Kidney organoid spheroids were micro-dissected and transferred to cell strainers in 6-well cell culture plate containing RB media one day before hypoxic treatment. The organoids were rinsed in DPBS and immediately fixed with 4% paraformaldehyde, followed by cryoprotection using a sucrose gradient (5% sucrose, overnight; 10%, 10 min; 13%, 10 min; 15%, 10 min; 20% 10 min). The organoids were embedded in OCT:20% sucrose (2:1) in a cryomold, followed by snap freezing on dry ice. 5 µM sections were rehydrated with PBS, permeabilized and blocked with 5% normal donkey serum in PBS containing 0.1% Triton X-100 for 60 min. The sections were incubated with primary antibodies diluted in 3% bovine serum albumin overnight at 4°C, followed by incubation with the appropriate secondary antibodies and mounting with Prolong Gold Antifade. Immunofluorescence images were captured using a 40x objective on a Leica DM IRB Inverted Fluorescence Microscope and then quantified using QuPath software^66^. For nuclear HIF1A protein expression, a total of 20 images were quantified and nuclear detection for each image was performed at a constant threshold. For nuclear ATF6 protein expression, a total of 30 tubules were quantified and nuclear detection from each tubule was performed at a constant threshold. For the quantitation of c-Cas3, ATP5A and pS6 protein expression, images containing 30-50 tubules were manually annotated using the closed polygon tool and fluorescence intensity was measured at a constant threshold using pixel classification for the specified protein channel. For COL1A1 and FN1 protein expression, a total of 20 images were quantified, and protein positive area was measured at a constant threshold using pixel classification for the specified protein channel.

### ELISA

Secreted VEGFA levels from kidney organoids were measured in organoid culture media in duplicate, using commercially available human VEGF ELISA kit following manufacturer’s protocol. Values were normalized to total protein content using Pierce BCA protein assay.

### Lipid droplet staining and quantification

Cryosectioned organoids mounted on slides were rehydrated with PBS three times for 5 min and permeabilized with 0.1% Triton X-100 in PBS for 60 min. The sections were incubated with the LipidSpot 488 at 1:100 dilution for 30 min, and rinsed once with PBS for 5 min, and mounted with Prolong Gold Antifade with DAPI. A total of 20 images were captured using a 63x oil objective. The droplet images were converted to grayscale, binarization by a constant threshold and quantified using Fiji software^66^. Cell counting from each cell image was performed at a constant threshold using QuPath. Based on these values, the number of lipid droplets per cell was calculated and results were plotted.

### Extracellular oxygen consumption assay

For the measurement of Oxygen Consumption Rate (OCR), protocol was applied as per manufacturer’s instructions. Briefly, 20 pre-picked organoids were added in wells pre-loaded with 150 µl of warm RB media, in triplicate. Then, 20 µl of reconstituted Extracellular O_2_ Consumption Reagent was added to each sample well. In parallel, 20 µl of culture media were added to blank wells. All wells were sealed with 2 drops of mineral oil and measured at 1.5 min intervals for 90-120 min at Ex/Em = 380/650nm. One experiment was performed and plotted ± SD.

### HK-2 cell culture and treatment

The immortalized human renal proximal tubular epithelial cell line, HK-2, was maintained in DMEM/F12 medium containing 5 mM glucose, prepared by mixing DMEM without glucose and Ham’s F12 Nutrient Mix at 1:1 ratio, supplemented with 10% Fetal Bovine Serum (FBS) and 1x antibiotic-antimycotic solution. Cells at passage 6 were seeded into 8-well chamber slides and cultured under normoxic conditions for 48h. Prior to hypoxic exposure, cells were pre-incubated for 30 min with two different doses of dapagliflozin,10 and 50 µM. Hypoxic culture (5% O_2_) was carried out for 24h using a Panasonic MCO-5M-PA multi-gas incubator, where O_2_ concentration was reduced by introducing nitrogen gas within 15-20 min of door closure.

### Measurement of ROS production in HK-2 cells

After 21h under hypoxia, 25 µM H2DCFDA, an indicator for ROS, was added to each well, mixed quickly and gently, and incubation was continued for an additional 3h under the same hypoxic conditions. Cell monolayers were then washed with PBS and fixed with 4% formaldehyde in PBS for 10 min. After another PBS wash, coverslips were mounted on the fixed monolayers using Fluoromount-G Mounting Medium, with DAPI. Slides were scanned using a ZEISS Axio Scan 7 microscope to capture blue (DAPI) and green (DCF) fluorescence. Automatic detection and quantification of green fluorescence were performed using QuPath. Each condition was assessed in triplicate, and three independent fields were analyzed for quantification where more than 300 cells were analyzed per condition. One experiment was performed and plotted ± SD.

### Bulk RNA-Sequencing and bioinformatic analysis

Total RNA from isolated organoid spheroids was prepared as detailed above. Library preparation using NEBNext Ultra II kit and sequencing using paired end read length of 150 bases on a NovaSeq4000 were performed by the University of Michigan Advanced Genomics Core. Reads were inspected for non-target sequences with FastqScreen^67^. Fastq read quality was determined using FastQC and reads aligned to the reference (ENSEMBL GRCh38.104) using STAR 2.7.8a^68^. Uniquely mapped reads were inspected for unusual distribution across known annotated features using Picard Tools. Gene level read counts were generated using HTSeq (version0.12.4)^69^ and normalized with voom^70^. The above tools, in conjunction with STAR read mapping statistics, Principal component analysis (PCA) and hierarchical clustering were used to identify and remove samples with abnormal expression profiles due to technical issues. Genes with ENSEMBL IDs mapping to a NCBI ENTREZ Gene ID were used for further analyses. DE genes between groups defined in each figure were extracted in the TIGR MultiExperiment Viewer application (unpaired analysis)^71^. Genes regulated with a false discovery rate (FDR) <0.05 were considered significant and used for further analyses with criteria defined in each figure.

### Single cell RNA-sequencing and bioinformatic analysis

Whole well organoid cultures were collected by scraping cells into ice-cold DPBS and dissociated with cold activate protease. Single cell suspensions were then submitted to the Advanced Genomics Core at the University of Michigan for library preparation and sequencing on a 10x Genomics Chromium Single Cell 3’ v3.1 platform. Raw sequencing data were processed with Cell Ranger for alignment to the human reference genome (ENSEMBLE GRCh38.104) and generation of feature-barcode matrices. Downstream analysis was performed using Seurat^72^. Cells expressing >500 genes were included in the analysis. Processing steps included log-normalization, scaling, identification of highly variable genes, dimensionality reduction with principal component analysis (PCA) and Uniform Manifold Approximation and Projection (UMAP), batch correction using the Harmony algorithm (v1.0)^73^, and unsupervised clustering at a resolution of 0.25.

### Canonical pathways, pathway-enrichment analyses, dot plots, volcano plots and heatmaps generation

QIAGEN Ingenuity Pathway Analysis software was used to identify regulated canonical pathways from DE genes. Canonical pathways with p-value<0.05 were considered significant. Dot plots were created using an in-house R script. The customized script used for creation of the dot plots is included with other analysis code in GitHub. MetaCore software was used to identify the “Process networks” enriched in the sets of differentially expressed proteins. Pathways with FDR <0.05 were considered significant. Heatmaps were generated using the Morpheus visualization software using the ‘transform values: subtract row mean, divide by row standard deviation’ option (which generates color intensity based on the Row Z-score). Pathways of interest were selected from the Pathway Browser of the Reactome Pathway Database^74^ and genes from each pathway were downloaded. Gene ontology (GO) molecular function was extracted from the PANTHER knowledgebase using EntrezGeneID as input. Volcano plots were generated using either GraphPad Prism or VolcaNoseR.

### Calculation of signature scores

The gene expression values for gene signatures were Z-transformed following a previously described calculation method^25,74^. In brief, for each gene, the expression was subtracted from the mean of all samples and divided by the standard deviation. A z-score was generated for each sample or patient. Statistical analysis of gene scores was performed using an unpaired parametric t-test with GraphPad Prism. P-values<0.05 were considered significant.

### Glucose flux and metabolomic analysis

One day prior to hypoxic treatment (D22), kidney organoids (15 pooled per condition) were micro-dissected and transferred to cell strainers placed in 6-well plates containing RB. On D23, kidney organoids were placed under hypoxia for 24h and were treated with ^13^C6-glucose (11 mM) in DMEM without phenol red for the last 3h of hypoxic treatment. Organoids were rinsed once with ice-cold ammonium acetate (50 mM) in LC-MS grade water, quenched with 200 µl of ice-cold LC-MS grade methanol, snap-frozen in liquid nitrogen, and stored at -80°C until further processing as previously described^75,76^. Briefly, the frozen organoids were sonicated on ice for 20 secs using an OmniRuptro 280 at 30% power and 30 pulse. Immediately after sonication, 200 µl of water and 400 µl of chloroform were added. The mixture was sonicated for 10 secs and placed on ice for 5 min. The samples were centrifuged at 15,000 x g for 10 min. The resulting upper layer (400 µl) was collected, dried and then reconstituted with 30 µl of LC-MS grade acetonitrile:water (2:1) and filtered before injecting 5 µl for analysis. Metabolites were analyzed on the Agilent 6456 quadrupole time-of-flight mass spectrometer coupled to Agilent 1290 LC as previously described^77^. Authentic standards of all measured metabolites were run separately and spiked into pooled samples for verification of metabolite identity and retention time.

### MALDI-MSI

Freshly harvested kidney organoids were embedded in a 7% gelatin mold using an isopentane/liquid nitrogen freezing method. For sectioning onto ITO-coated slides, gelatin-embedded organoids were sectioned at 10 μm thickness using a Leica CM1950 Cryostat. The matrix application was performed using the automated HTX M3+ sprayer. 1,5-diaminonaphthalene (DAN, 5.5 mg/ml in 50% ethanol and 5% HCl) was used for negative ion mode and or 2,5-dihydroxybenzoic acid (DHB, 40 mg/mL in 50% methanol) was used for positive ion mode. MALDI-MSI was conducted using Q Exactive HF-X hybrid quadrupole-Orbitrap mass spectrometer coupled with an elevated pressure MALDI/ESI interface, following our previously described protocol^78^. Mass spectra were acquired at a spatial resolution of 20 μm over an m/z range of 100–1000 in both polarities, with a resolving power of 120,000 at m/z 200. Metabolite annotation was performed using METASPACE^79^. Pre-MALDI optical images were used to assess organoid quality and to select regions of interest (ROIs) in METASPACE for downloading ion intensities of annotated molecules at 20% FDR using CoreMetabolome, HMDB^80^, KEGG, and SwissLipids^81^ databases.

### Metabolite quantification by LC-MS/MS

Amino acids and related metabolites were quantified in organoid culture supernatant using a targeted liquid chromatography mass spectrometry (LC-MS) approach. For sample preparation, 10 µl of supernatant was transferred into a 96-well plate and extracted with pre-chilled 80% MS grade methanol and stored at -20°C for 30 minutes. Proteins were precipitated and 50 µl of the supernatant was carefully transferred to a new 96-well plate for LC-MS analysis. For amino acids and related metabolites, chromatographic and mass spectrometric analyses were performed on a Q Exactive HF-X Orbitrap MS equipped with a heated electrospray ionization (HESI) source operated in positive mode and interfaced with a Thermo Vanquish UHPLC system. A 2.5 µl aliquot of each sample was injected via autosampler. For amino acid and related metabolite separation, hydrophilic interaction chromatography (HILIC) was employed using an Agilent ZORBAX HILIC PLUS column (2.1 × 100 mm, 3.5 µm particle size). The mobile phase consisted of 10 mM ammonium formate with 0.05% formic acid (FA) in water (Solvent A) and 0.05% FA in acetonitrile (Solvent B). The flow rate was set at 0.3 ml/min (PMID: 37752301). For fatty acids analysis, chromatographic separation was performed on a Waters XBridge Peptide BEH C18 column (2.1 × 100 mm, 2.5 µm particle size) with 10 mM ammonium bicarbonate and 0.05% FA in water (Solvent A), and 0.05% FA in methanol (Solvent B), at a flow rate of 0.25 ml/min with mass spectrometry data acquired in negative ionization mode. High-resolution MS data were acquired in full scan and data-dependent MS/MS modes. Data processing, including peak integration and quantification, was performed using Thermo Xcalibur Quant Browser software. Metabolite identification was confirmed using retention time of standards, exact mass (within ± 5 ppm), and MS/MS spectral match to authentic standards.

### Proteomic Sample Preparation

Organoids (N = 12 per sample) were resuspended in 150 µl of 4% SDS buffer containing 100 mM HEPES (pH 7.4), 10 µg/ml DNase I, protease and phosphatase inhibitors. Protein extraction was assisted by 15 seconds of bead beating using Storm Pro Bullet blender (2 steel beads per sample, amplitude 8). Following centrifugation at 9,000 x g for 3 min, the samples were heated at 95°C for 5 min. An aliquot was taken for protein quantification using the BCA assay. Proteins were reduced and alkylated by adding 5 mM (Tris(2-carboxyethyl) phosphine) (TCEP) and 10 mM of Chloroacetamide (CAA), followed by heating at 95°C for 5 min. Samples were stored at −18°C. For cell culture supernatants, samples were lysed as the spheroids but with a lower buffer concentration: 2% SDS buffer containing 50 mM HEPES (pH 7.4) and 5 µg/mL DNase I. Protein cleanup was performed using the Single-Pot Solid-Phase-enhanced Sample Preparation (SP3) method^82^. Briefly, protein lysates were mixed with Magnetic Carboxylate SpeedBeads 50:50 hydrophilic:hydrophobic at a 1:10 (v/v) bead-to-sample ratio in the presence of 80% ethanol and incubated at room temperature for 18 min. Beads were captured using a magnetic rack, and the supernatant was discarded. The beads were washed twice with 90% acetonitrile to remove contaminants. After drying, beads were resuspended in 50 mM Triethylammonium bicarbonate buffer (TEAB, pH 8.0), and proteins were digested overnight at 37°C with trypsin at an enzyme-to-protein ratio of 1:50 under constant shaking. Tryptic peptides were dried and purified using C18 StageTips conditioned with 100 µl of methanol and equilibrated twice with 200 µl of 0.1% FA. Acidified peptide samples (final FA concentration 0.5%) were loaded onto the tips by centrifugation at 1,000 × g. Bound peptides were washed with 0.1% FA and eluted with 60% acetonitrile in 0.1% FA. Eluates were vacuum-dried and resuspended in 0.1% FA for LC-MS/MS analysis.

### Proteomic LC-MS/MS analysis

Tryptic peptides in 0.1% FA were analyzed using an UltiMate3000 nanoHPLC system coupled to an Orbitrap Exploris 480 mass spectrometer equipped with a high-field asymmetric waveform ion mobility spectrometry (FAIMS) Pro interface. Peptide separation was performed using a two-column setup, consisting of a trap column (Acclaim PepMap 100 C18, 3 µm particle size, 2 cm x 75 µm) and an analytical column (Aurora Ultimate 25 cm x 75 µm, C18) operated at 400 nl/min. The analytical column was maintained at 50°C throughout the 120 min gradient. Peptides were separated using a binary gradient with solvent A (H2O + 0.1% FA) and solvent B (acetonitrile + 0.1% FA). The gradient began at 2% B, increasing linearly to 8% B over the first 10 min, continued to 25% B by 90 min, then ramped to 35% B at 100 min, and further increased to 90% B between 100 and 109 min. After a 1 min hold at 90% B, the column was re-equilibrated to 2% B from 110 to 120 min. The mass spectrometer was operated in positive ion mode with a static spray voltage of 1800 V, an ion transfer tube temperature of 275°C, and a total carrier gas flow of 3.8 l/min. The RF lens was set to 40% and the default charge state was set to 3. The FAIMS Pro interface was run at standard resolution, alternating between two compensation voltages (CVs): 45 V and 65 V in separate DIA scan cycles. MS1 full scans were acquired in the Orbitrap with a resolution of 120,000, scanning from m/z 380-1500, using 1 microscan, automatic injection time, and a custom automatic gain control (AGC) target of 300%. DIA MS2 scans were acquired at a resolution of 30,000 using 40 custom isolation windows (15 m/z each) spanning 4001000 m/z, with HCD fragmentation at 28% normalized collision energy, automatic injection time, and a normalized AGC target of 1000%. All scans were recorded in profile mode, with a 3 sec cycle time per CV. Sample injection was performed using microliter pickup mode. The sample compartment was maintained at 8°C, and pressure, flow, and temperature parameters were monitored continuously throughout the run.

### Proteomic data analysis

Raw data were analyzed using Spectronaut software using the canonical human reference proteome from UniProt^83^ (UP000005640_9606, 20,596 entries, January 2024) for the Pulsar database search. Identification was performed using the “directDIA+ (Deep)” workflow with default parameters. Enzyme specificity was set to Trypsin/P, carbamidomethylation of cysteine was used as a fixed modification (+57.021464), while oxidation of methionine (+15.994914) and protein N-terminal acetylation (+42.010565) were set as variable modifications. For DIA analysis, non-linear iRT-to-RT regression was applied. Precursor-level FDR was controlled at 1%, and protein-level FDR was set to 1% at the experiment level. Protein inference was based on the IDPicker algorithm. Quantification was performed at the MS2 level using area under the curve with the MaxLFQ algorithm as implemented in Spectronaut. Cross-run normalization was enabled, and no imputation strategy was applied. The spectronaut output (normalized Log_2_ quantities) were further processed using the DEP2 package^84^ in R and RStudio, which internally uses limma^85^ for comparison testing. We applied a threshold for significance based on Benjamini-Hochberg adjusted or unadjusted p-value≤0.05 and an absolute Log_2_ FC≥0.58.

### Kidney pathology

Diagnosis in NEPTUNE was assigned by study pathologists based on biopsy reports and/or digital images. The degree of acute tubular injury (ATI) was visually assessed, scored by 2 to 5 pathologists using biopsy whole slide images of stained sections, recorded as the percent of cortex involved, and averaged across the pathologists’ measures^26,27^.

### Quantification and Statistical Analysis

Statistical analysis was performed using the tests as indicated within figure legends. Data from at least three independent experiments were plotted as mean ± standard error of the mean (SEM), unless otherwise stated, using GraphPad Prism 10 software. Groups were compared using unpaired t-test (two-tailed), as appropriate. Welch’s correction was used for groups with unequal standard deviations in unpaired Student’s t-test. One-way Anova was used to compare three or more independent groups, whereas Two-way Anova was used for groups with 2 independent variables. Statistical significance was displayed as an asterisk in corresponding graphs and was denoted as follows: ∗p<0.05, ∗∗p<0.01, ∗∗∗p<0.001.

