## Supplementary Figures S1-S6 for "Dapagliflozin mitigates hypoxia-induced metabolic stress, kidney tubular cell death and fibrosis"

#### Figure S1. hPSC-kidney organoids exhibit expected transcriptional and metabolic hypoxic response

**(A)** Mean oxygen levels inside the closed hypoxia chamber measured at different time points throughout the hypoxia experiment.
**(B)** Immunofluorescence showing HIF-1α expression in T1D-derived organoids (49E, male and 51A, female), after 24h normoxia (top) or hypoxia (bottom).
**(C)** Expression of *EGLN1*, *SLC2A1* and *VEGFA* transcripts in T1D-49E organoids after 24h hypoxia (n=1 +/- SD, technical duplicates).
**(D)** Immunofluorescence comparing ATP5A expression in hypoxic and normoxic organoids, with quantification (n=3).
**(E)** Scatter plot comparing oxygen consumption levels in hypoxic and normoxic organoids (n=1 +/- SD, technical triplicates).
**(F)** Schematic of Hypoxia protocol #1b which includes 1 day of hypoxia followed by 2 days of reoxygenation. Black arrowheads indicate sample collection times at D24, 25 and 26 (0, 24 and 48h, post-reoxygenation, respectively).
**(G)** Graph comparing culture media glucose levels of organoids at 0, 24 or 48h post-reoxygenation (n=1 +/- SD, technical duplicates).
**(H)** Expression of *SLC2A1*, *VEGFA*, *TGFB1* and *COL1A1* transcripts in organoids at 0, 24 or 48h post-reoxygenation (n=3).
**(I-K)** Immunofluorescence comparing HIF-1α (I), GLUT-1 (J) and PODXL (K) expression in hypoxic following 2 days of reoxygenation vs. normoxic organoids at D26.

Data displayed as mean +/- SEM; n=biological replicates; Unpaired t-test (C, D), Two-way Anova (G, H); *$p<0.05$, **$p<0.01$, ***$p<0.001$, ns, not significant. IF markers: Nuclei (DAPI), tubular epithelial membrane (N-cadherin/pan-CK), Podocytes (Podocalyxin (PODXL)); white scale bars, as indicated.

**Related data** Figure 1.

#### Figure S2. An organoid-based transcriptional signature of hypoxia identifies individuals with tubular injury and poor kidney disease outcome

**(A and B)** Summary expression of 112-unique genes of Curated HIF (A) and 334-unique genes of Hypoxic Organoid (B) Signatures (in Fig. 2A) in kidney tissue transcriptome clusters (in Fig. 2C). Boxplots represent median (white line), interquartile range (box), and minimum and maximum values (whiskers).
**(C)** Table showing NEPTUNE tubular morphological features used for scoring Acute Tubular Injury (ATI) recorded as the percent of cortex (0–100%) involved by injury.
**(D and E)** Boxplots showing percentage of interstitial fibrosis (D) and tubular atrophy (E) based on ATI scoring of human tissue clusters. Boxplots represent median (black line), interquartile range (colored box), and minimum and maximum values (whiskers).

**(F)** Scatter plot for HIF1A transcript levels based on ATI scoring of human tissue clusters. Median expression, black line.
**(G and H)** Dot plot of individual tissue transcriptome summary gene expression of 112-unique genes of Curated HIF (G) and 334-unique genes of Hypoxic Organoid (H) Signatures (color corresponds to cluster in (A and B)), sorted by corresponding tissue ATI score. Median expression, black line. Statistical comparisons between ATI scores shown; *$p<0.05$, **$p<0.01$, ***$p<0.001$.
**(I)** Bubble plot of gene contribution of Signature gene sets (in Fig. 2A) to Reactome pathways, represented as percent of genes in each of the Signatures.
**(J)** Pie chart of GO Molecular Functions from the Panther Classification System represented by the 94 genes, not expressed in human kidney tissue (from Fig. 2D), of the 377-gene Hypoxic Organoid Signature.

**Related data** Figure 2, Table ST4-5.

**Figure S3. Hypoxia induces tubular epithelial cell apoptosis, alters cell energetics and enhances fibrosis**

**(A)** Immunofluorescence image showing QuPath analysis for cleaved caspase 3 expression (green spots) in organoid tubular cells, delineating tubular area in blue (white arrows) and excluding lumen area in pink (yellow arrows).
**(B)** Graph comparing culture media glucose levels of organoids after 24 or 48h hypoxia versus normoxia (n=1 +/- SD, technical triplicates).
**(C)** Expression of *HAVCR1* transcript in organoids after hypoxia for 24h at D24 (left, n=3), 48h at D24 (middle, n=3), or 48h at D20 (right, n=2).
**(D)** Immunofluorescence comparing KIM1 expression in normoxic versus 24h hypoxic organoids.
**(E)** Graph comparing bulk transcriptional expression of genes from the 377-gene Hypoxic Organoid Signature in organoids at D20 post-hypoxia or D25 post-reoxygenation, versus normoxia.
**(F)** Immunofluorescence and quantification (n=3) of FN1 expression in D25 normoxic (top) versus reoxygenated (bottom) organoids.
**(G)** Bar plot comparing the expression of 6 genes of interest at the RNA, tissue protein and secreted protein levels in organoids at D20 post-hypoxia (top) or D25 post-reoxygenation(bottom), versus normoxia.

Data displayed as mean +/- SEM; n=biological replicates; Two-way Anova (B), unpaired t-test (C, E); *$p<0.05$, ***$p<0.001$, ns, not significant. IF markers: Nuclei (DAPI), tubular epithelial membrane (pan-CK); white scale bars, as indicated.

**Related data** Figure 3

**Figure S4 Hypoxia stimulates unfolded protein response and suppresses expression of mitochondrial β-oxidation proteins**

**(A)** UMAP clustering (14 clusters) of 38,100 cells as profiled by scRNA-seq of organoids under normoxia or hypoxia, treated or not with vehicle control (VC) or dapagliflozin for 24h.
**(B)** Dot plot showing expression of main cell marker genes in clusters identified in (A).
**(C)** Dot plot comparing expression of key UPR genes in kidney-specific cell clusters of normoxic versus hypoxic organoids.
**(D and E)** Graph comparing *sXBP1* to *tXPB1* transcript expression in D20 acutely hypoxic (D, n=3) or D25 reoxygenated (E, n=3), versus normoxic organoids.
**(F)** Immunofluorescence comparing ATF6 expression in organoids after 48h hypoxia versus normoxia.

Data displayed as mean +/- SEM; n=biological replicates; Welch's t test (D), unpaired t-test (E); *$p<0.05$, ns, not significant. IF markers: Nuclei (DAPI); white scale bar, as indicated.

**Related data** Figure 4

**Figure S5. Dapagliflozin protects kidney tubular epithelial cells from hypoxia-mediated injury via decrease in reactive oxygen species**

**(A)** Dot plot of transcriptional expression of *SLC5A2 and SLC2A1/2/3/4* in tubular cells under normoxia or hypoxia +/- dapagliflozin for 24h, from single cell profiled organoids.
**(B and C)** Immunofluorescence comparing GLUT-1 (B) and HIF-1α (C) expression in hypoxic organoids treated with VC (top) or dapagliflozin (bottom) for 24h.
**(D)** Expression of *TGFB1* and *HAVCR1* transcripts in hypoxic organoids +/- dapagliflozin for 48h (n=3).
**(E)** Expression of *sXBP1* transcript in D20 acutely hypoxic (left) or D25 reoxygenated (right) organoids, +/- dapagliflozin (n=3).
**(F)** Immunofluorescence quantification (n=3) of nuclear ATF6 expression in tubules of hypoxic organoids +/- dapagliflozin for 24h.
**(G)** Stacked bar plots comparing the effect of dapagliflozin on differential expression of glycolysis proteins in D20 post-hypoxic (top graph) or D25 post-reoxygenated (bottom graph) organoids (orange bars), to the effect of hypoxia on differential expression of proteins relative to normoxia (blue bars) at the same time points.
**(H)** Stacked bar plot comparing the effect of dapagliflozin on differential expression of UPR proteins in D25 post-reoxygenated organoids (orange bars) to the effect of hypoxia on differential expression of proteins relative to normoxia (blue bars) at the same time point.
**(I)** Volcano plots of DE proteins in organoids secreted compartment at D20 post-hypoxia (top) or D25 post-reoxygenation (bottom) +/- dapagliflozin.

Data displayed as mean +/- SEM; n=biological replicates; Two-way Anova (D), unpaired t-test (E, F); ns, not significant. IF markers: Nuclei (DAPI), tubular epithelial membrane (N-cadherin/Pan-CK); white scale bars, as indicated.

**Related data** Figure 5.

**Figure S6. Integrated transcriptomic and proteomic analysis reveals markers of dapagliflozin-mediated modulation of hypoxia-induced cellular stress response**

**(A)** UMAP clustering (11 clusters) of 10,661 cells profiled by scRNA-seq of D25 reoxygenated organoids +/- dapagliflozin for 24h (top) and corresponding dot plot (bottom) showing expression of cell type-specific genes in identified clusters.
**(B)** Graph comparing expression of genes, from the 377-gene Hypoxic Organoid Signature, in proximal tubular cells at 24h post-hypoxia (left) or 5 days post-reoxygenation (right), +/- dapagliflozin.
**(C)** Graph comparing expression of genes, from the 377-gene Hypoxic Organoid Signature, in bulk transcriptional profiling of D20 acutely hypoxic or D25 reoxygenated organoids +/ dapagliflozin.
**(D)** Graph comparing expression of genes, from the 61 DE proteins with dapagliflozin treatment, in bulk transcriptional profiling of D20 acutely hypoxic or D25 reoxygenated organoids +/ dapagliflozin.
**(E)** Graphs showing correlation between RNA and protein expression levels in organoid secreted and tissue compartments, at D20 post-hypoxia (top graph) and D25 post-reoxygenation (bottom graph), +/- dapagliflozin.
**(F)** Volcano plot comparing dapagliflozin's effect on differential protein expression of 43-gene Overlap Signature set in organoid tissue compartment at D20 post-hypoxia (top) and D25 post-reoxygenation (bottom).
**(G)** Dot plots comparing dapagliflozin's effect on hypoxic organoid cell transcriptional expression of genes of interest (from Fig. 6F), post-hypoxia (top plot) and post-reoxygenation (bottom plot).
**(H)** UMAP clustering (11 clusters) of human kidney tubular cell types from healthy or proteinuric individuals (FSGS and MCD) profiled by sc/snRNA-seq. UMAP on the right shows relative sc to sn contribution to the dataset.

**Related data** Figure 6.

Supplementary Figure S1

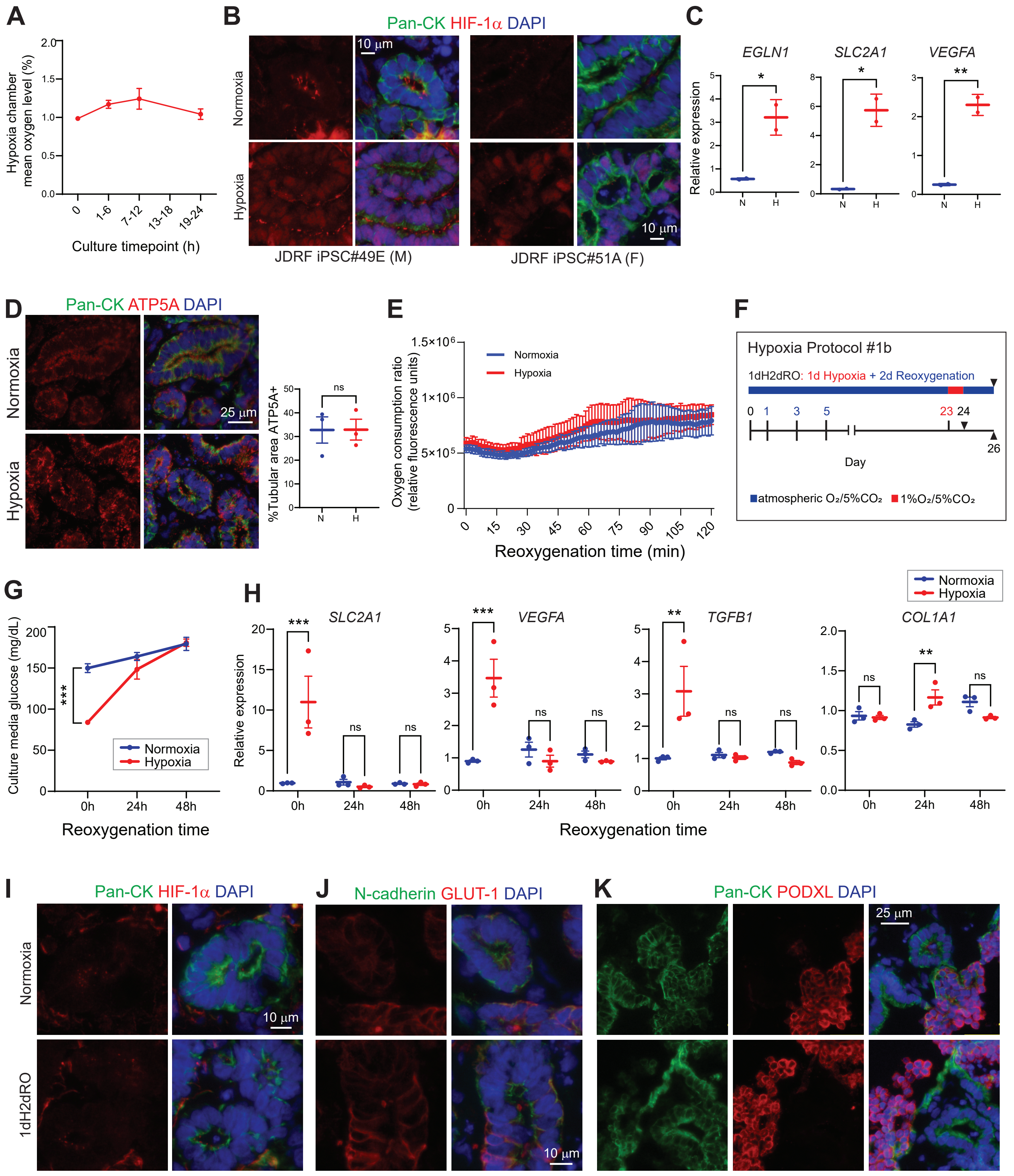

Supplementary Figure S2

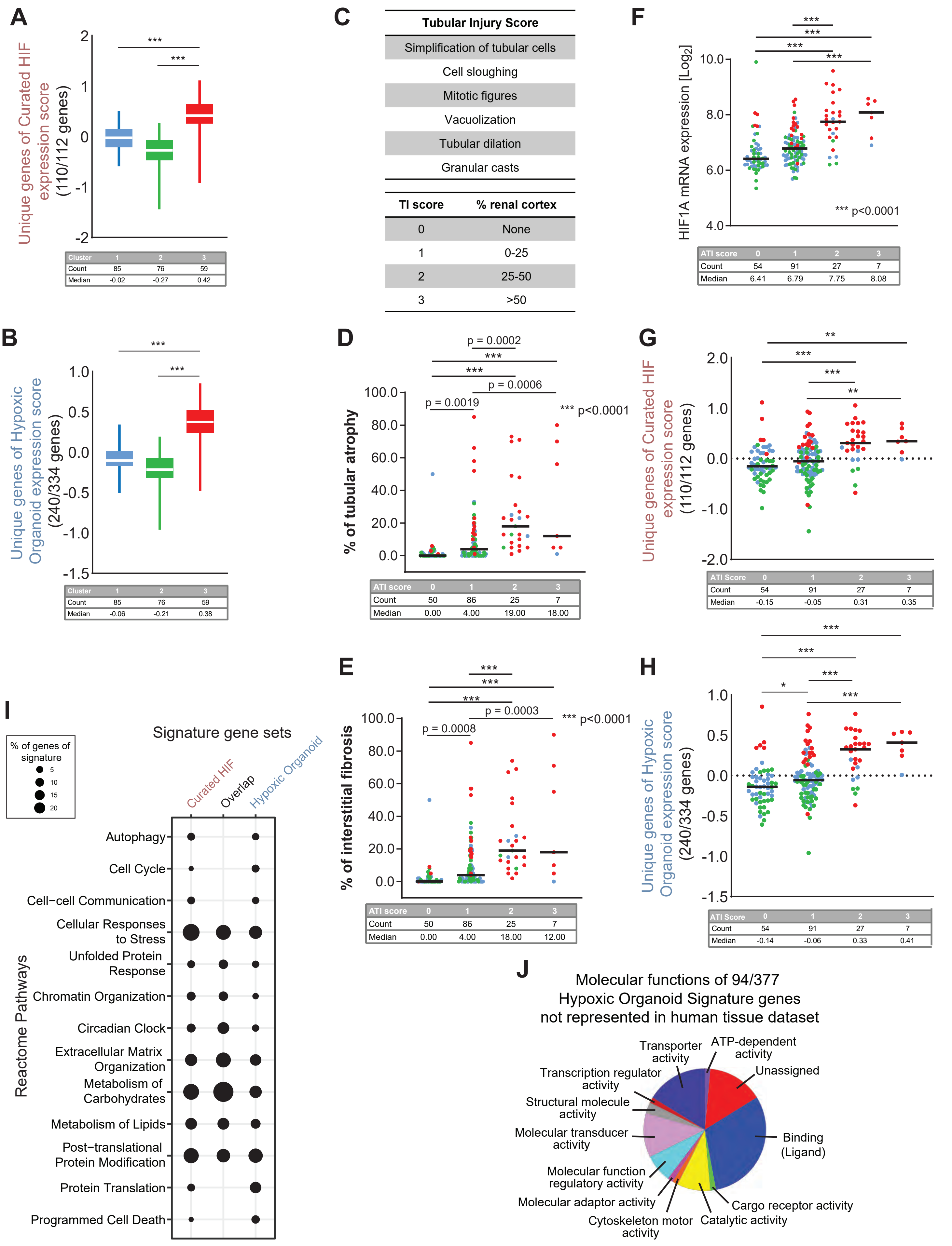

Supplementary Figure S3

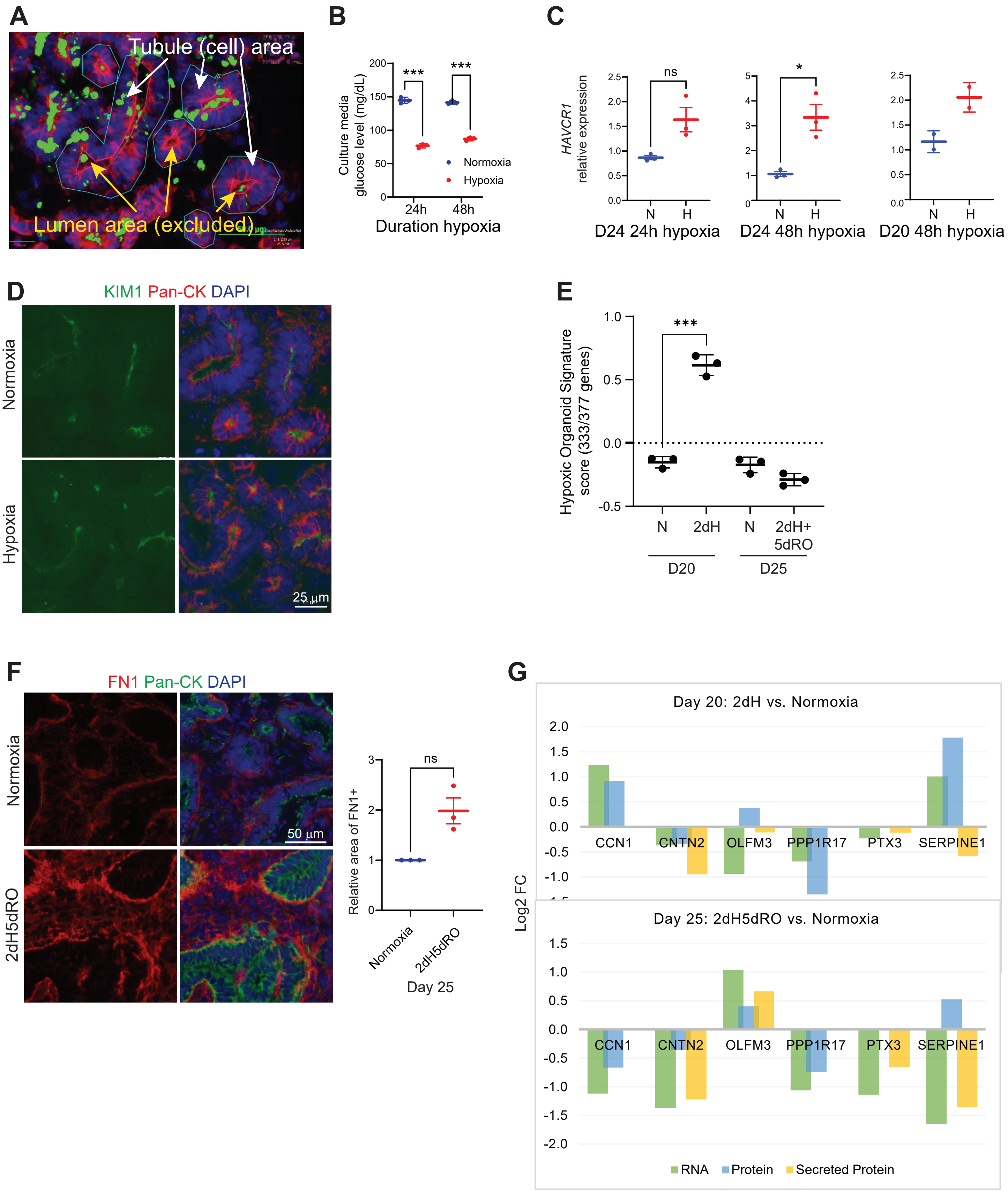

Supplementary Figure S4

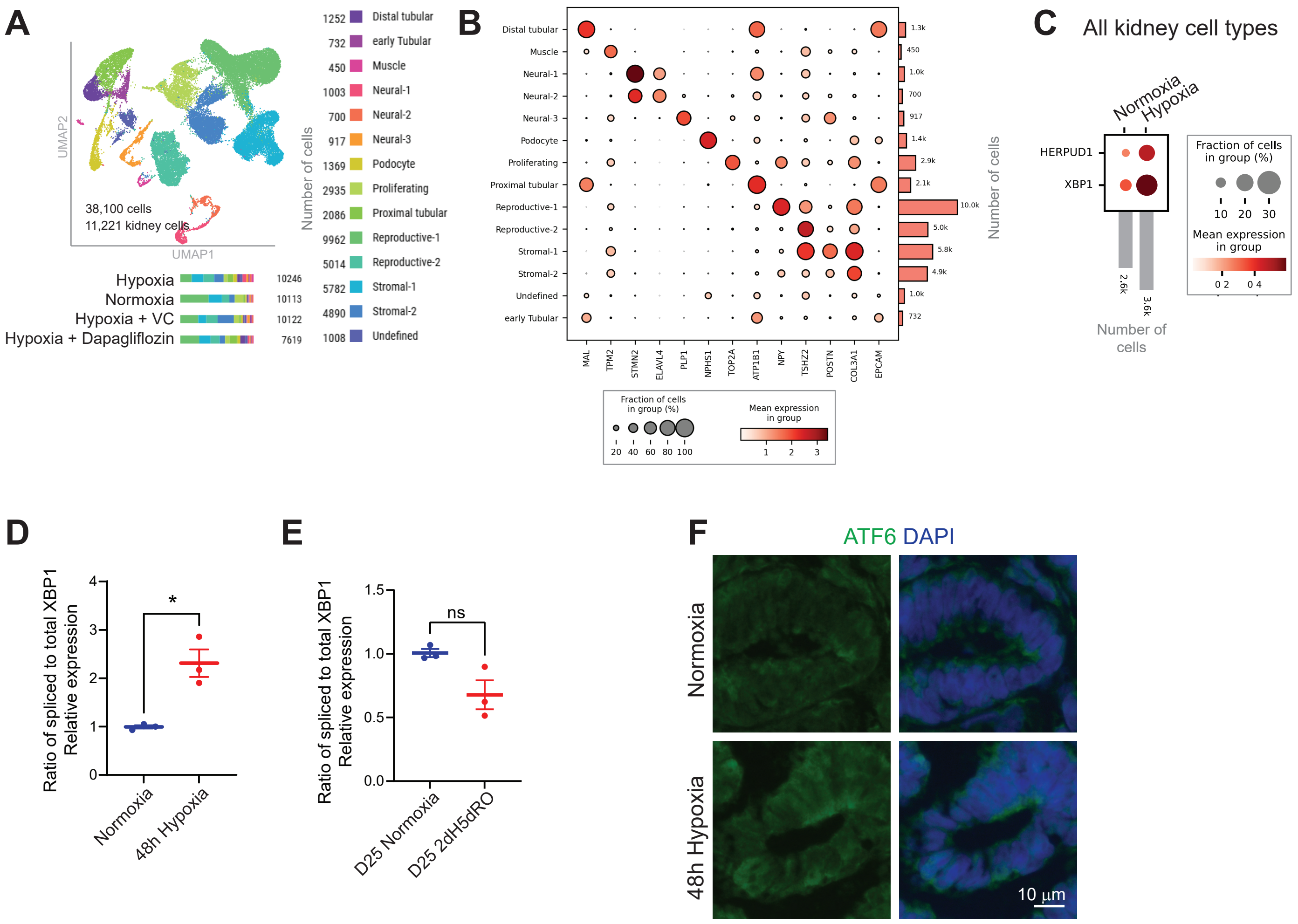

Supplementary Figure S5

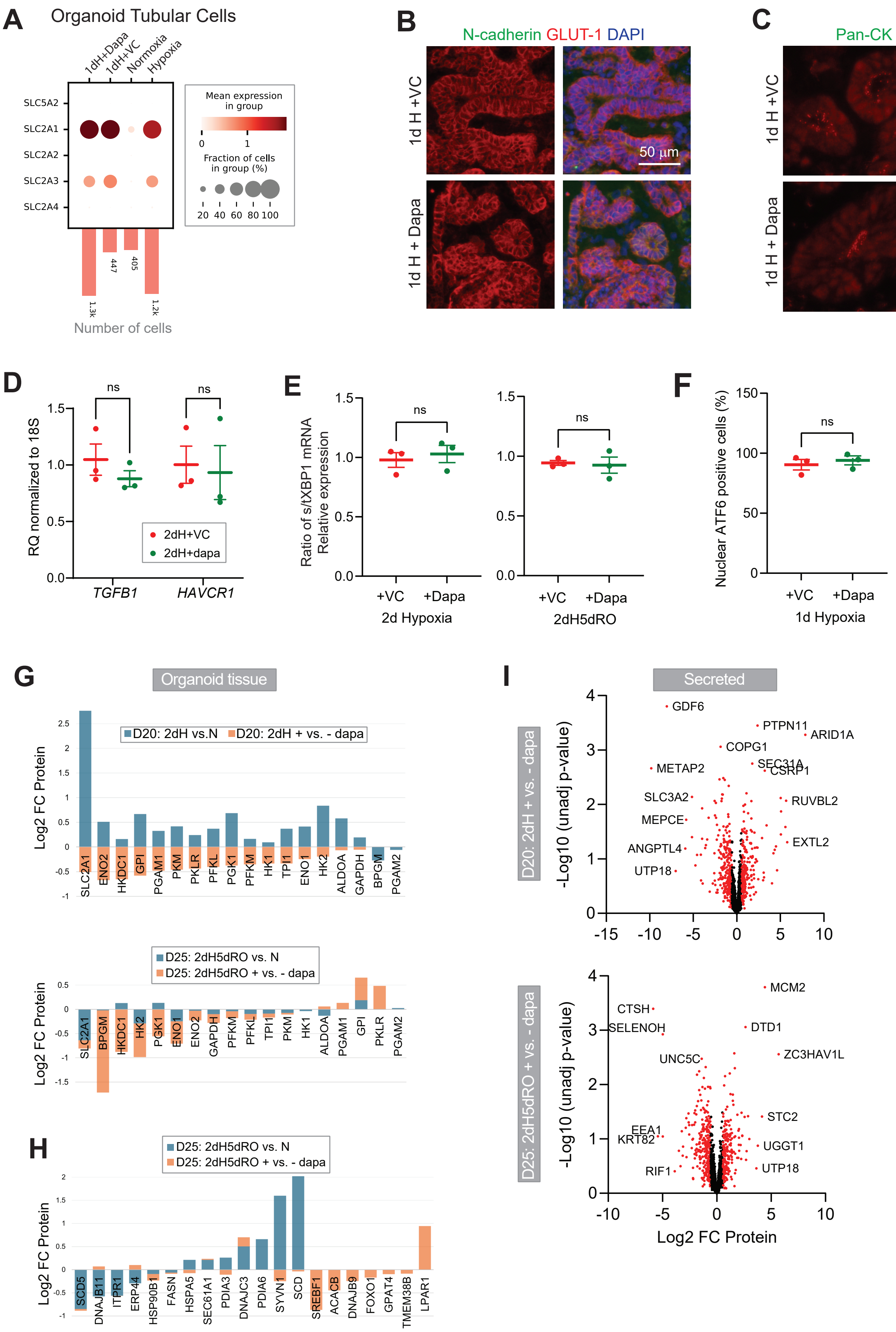

### Supplementary Figure S6

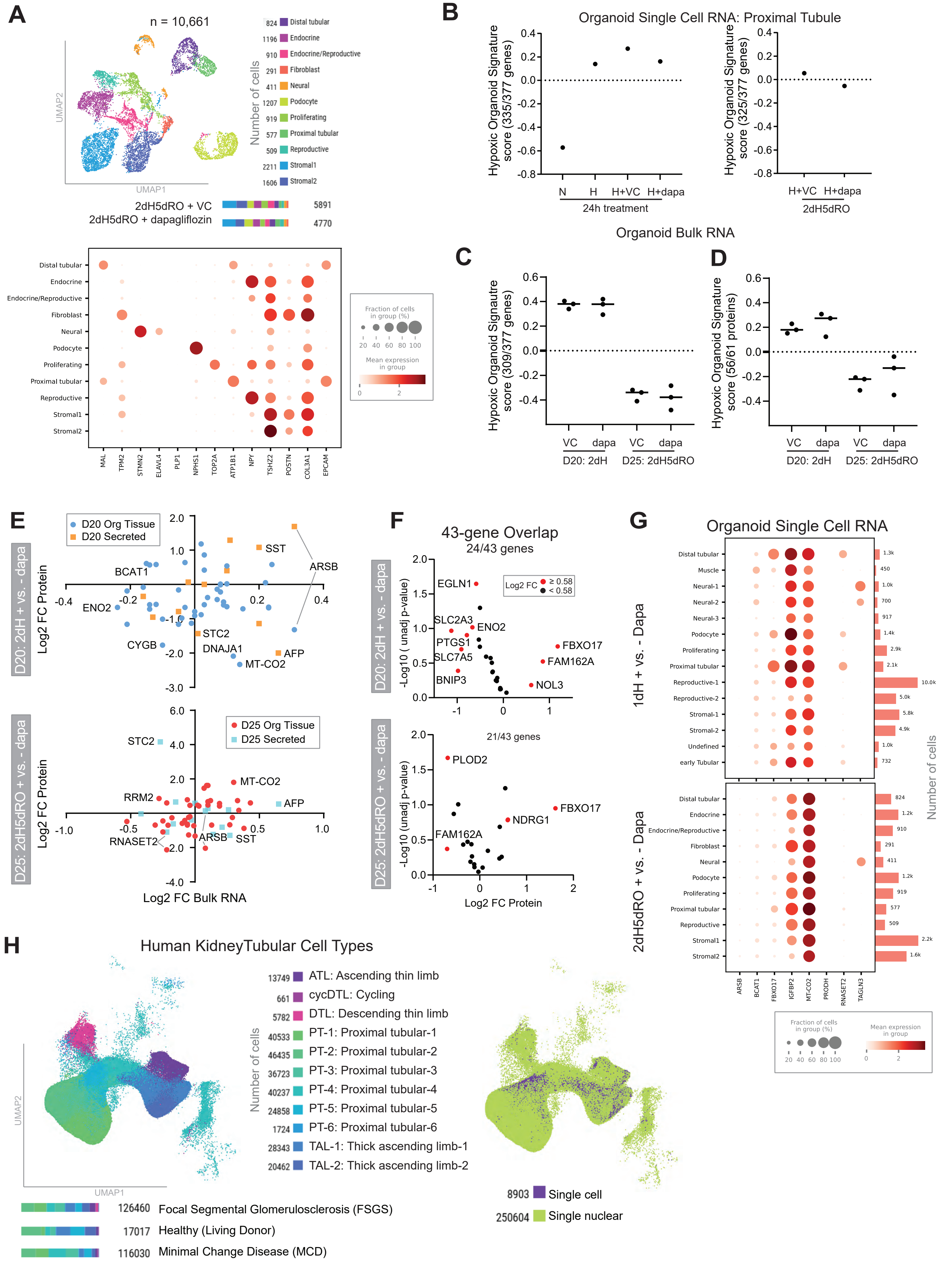
